# Plasma cell identity escape drives resistance to anti-BCMA T-cell–redirecting therapy in multiple myeloma

**DOI:** 10.64898/2025.12.08.692978

**Authors:** Francesco Maura, Ciara L. Freeman, Kylee H Maclachlan, Marta Larrayoz Ilundain, Bachisio Ziccheddu, Erin W Meermeier, Juan-Jose Garces, Michael Durante, Holly Lee, Ross Firestone, Meghan Menges, Kayla Reid, Benjamin Diamond, Marios Papadimitriou, Ariosto Siqueira Silva, Praneeth Reddy Sudalagunta, Noemie Leblay, Sungwoo Ahn, Ethan Fuller, Edward Briercheck, Meaghen E Sharik, Megan T Du, Phaedra Agius, Doris K Hansen, Xiaofei Song, Xiaohong Zhao, Mark B Meads, Jamie Teer, Tomas Jelinek, Patrick Hagner, Rachid Baz, Melissa Alsina, Anusuya M. Ramasubramanian, Eric Smith, Sergio A. Giralt, Sham Mailankody, Leif P. Bergsagel, Paola Neri, Marta Chesi, Jose A. Martinez-Climent, Frederick L. Locke, Nizar J. Bahlis, Ola Landgren, Issam Hamadeh, Saad Z. Usmani, Kenneth H. Shain

**Affiliations:** Myeloma Service, Memorial Sloan Kettering Cancer Center, New York, NY, USA; Myeloma Service, Sylvester Comprehensive Cancer Center, University of Miami, Miami, FL, USA; Moffitt Cancer Center, Tampa, FL, USA; Department of Hematology, Center for Applied Medical Research, Cancer Center Clinica Universidad de Navarra, IDISNA, CIBERONC, University of Navarra, Pamplona, Spain; Mayo Clinic, Scottsdale, AZ, USA; Arnie Charbonneau Cancer Institute, University of Calgary, Calgary, Canada; Dana Farber Cancer Institute, Boston, USA; Department of Haemato-oncology, University Hospital Ostrava and Faculty of Medicine, University of Ostrava, Ostrava, Czech Republic; Executive Director, Translational Development, Bristol-Myers Squibb, Summit, NJ 07901

## Abstract

Chimeric antigen receptor T-cell (CART) and T-cell engager (TCE) therapies targeting B-cell maturation antigen (BCMA) are transforming the treatment landscape for relapsed multiple myeloma (MM). However, despite impressive initial response rates, most patients eventually relapse. To investigate this unmet medical need, we applied whole-genome sequencing (WGS) to MM cells from cohorts of 102 relapsed patients treated with anti-BCMA CART and TCE therapies. Several genomic alterations were associated with clinical outcomes, particularly primary refractoriness, including high genomic complexity and mutations in genes regulating plasma cell identity, which predicted resistance to therapy. Single-cell RNA sequencing further revealed that MM cells from refractory patients exhibited high proliferation signatures and reduced expression of *TNFRSF17* (encoding BCMA), while were less enriched for plasma cell–associated transcriptional programs, a phenomenon we term “plasma cell identity escape.” This profile was strongly associated with immune dysregulation of CD8 T cells including increased activation and exhaustion. This evolution of MM toward a more proliferative and lineage-divergent state, refractory to the anti-BCMA T-cell redirecting therapies, was functionally validated in preclinical MM mouse models. Collectively, our results comprehensively define the cellular and molecular mechanisms underlying primary resistance to anti-BCMA therapies.

## INTRODUCTION

Chimeric antigen receptor T cells (CART) and bispecific T-cell engagers (TCE) have revolutionized the treatment landscape in patients with relapsed/refractory multiple myeloma (RRMM).^1^ In recent years, two different anti-BCMA CART products have been approved by the FDA after showing an overall response rate (ORR) >70%: Idecabtagene vicleucel (ide-cel) and Ciltacabtagene autoleucel (cilta-cel).^2–4^ Most recently, three TCEs directed against BCMA (Teclistamab, Elranatamab, and Linvoseltamab) obtained FDA approval as well, having showed an ORR of 61-63% in RRMM.^5–7^ ^8^ However, in stark contrast to diffuse large B-cell lymphomas (DLBCL) and B-cell leukemias,^9,10^ none of these therapies have proven curative for RRMM, and most patients inevitably relapse.^2,3,5,6,11,12^

RRMM patients treated with anti-BCMA T-cell redirecting therapies can be broadly categorized into three groups: i) long responders who achieve durable remissions, ii) patients who relapse after an initial response lasting more than six months, and iii) refractory patients who either fail to respond or relapse within the first six months. Recent studies indicate that progression after initial response to TCE is frequently driven by the acquisition of non-truncating missense mutations or in-frame deletions in the extracellular domain of BCMA, which abrogate the efficacy of TCE, such as Teclistamab and Elranatamab, despite detectable surface BCMA expression. ^13,14^ These mutations typically arise 6–9 months into TCE-based therapies and account for 50–60% of relapses in this group,^15^ whereas such alterations are not observed after CART-cell therapy.^16^ While progression after an initial response to both CART and TCE therapies can be driven by biallelic loss of *TNFRSF17*, this mechanism is relatively uncommon, occurring in only ∼5% of patients.^17–19^ High tumor burden is another factor associated with refractory disease and immune dysfunctions, but its assessment is complicated by the inherent heterogeneity of RRMM clinical presentations.^20,21^ Moreover, multiple lines of therapy can drive evolution toward more aggressive disease, including extramedullary disease (EMD), which is often associated with primary refractoriness.^5,20–23^ Despite these insights, robust biomarkers capable of predicting patient responses to anti-BCMA CART or TCE therapies remain lacking, limiting our ability to identify high-risk patients and to tailor alternative treatment strategies.

To define the key genomic determinants of response to anti-BCMA immunotherapies, we analyzed whole-genome sequencing (WGS) and whole-exome sequencing (WES) from 114 samples collected from 102 patients with RRMM treated with either anti-BCMA CART or anti-BCMA TCEs. Our analyses revealed that patients who were refractory to anti-BCMA CART or TCE therapies shared distinct genomic features, including high genomic complexity and loss of genes involved in plasma cell identity. These alterations were associated with specific single-cell RNA (scRNAseq) expression signatures characterized by proliferative programs, genomic instability, reduced plasma cell differentiation and increased T CD8 exhaustion/activation. This distinct profile was named “*plasma cell identity escape*”. Importantly, this entity underlying refractoriness to CART and TCE therapies was validated in two distinct MM mouse models with different genomic and transcriptomic backgrounds treated with anti-BCMA TCEs. Defining the existence of this plasma cell identity escape phenomenon is clinically relevant, as it provides prognostic information beyond standard clinical features and enables the identification of patients unlikely to benefit from anti-BCMA immunotherapies, for whom alternative therapeutic strategies are warranted.

## RESULTS

### Anti-BCMA CART and TCE cohorts

To investigate the genomic mechanisms involved in primary refractoriness and resistance to anti-BCMA immunotherapies, we interrogated 114 WGS (60X median coverage) and 10 WES generated from a total of 102 patients treated with either anti-BCMA CART (n=77) and/or anti-BCMA TCE (n=26). One patient was treated with CART and TCE with a 10-month interval between the two treatments, and the same sample was used for both groups. Overall, all pre-treatment samples were collected within a year from the anti-BCMA therapy. Two patients treated with TCE underwent WGS at progression after one month; they were included in the subsequent clinical analysis, as clonal evolution is generally not observed within such a short timeframe. ^24,25^ Among the CART treated patients, 58 (74%) and 16 (26%) were treated with ide-cel and cilta-cel. A total of 20 WGS were generated from samples collected at the time of progression after CART and TCE, 17 of which had both time points available. Three patients with only one sample collected at progression were treated with Orvacabtagene autoleucel (orva-cel), ide-cel and cilta-cel. Three patients had WGS performed on samples collected at day +30, prior to overt disease progression, but with sufficient plasma cell involvement to yield high-quality WGS results. Among patients treated with anti-BCMA TCE (n=26), WGS was available for 24 samples collected prior to treatment and 8 samples collected following anti-BCMA TCE therapy (n=2 Teclistamab and n=6 Elranatanab); 6 of these patients had samples and WGS at both time points. Key clinical and demographic information are summarized in **Supplementary Table 1-2**. The CART–treated patients had a median progression-free survival (PFS) of 13 months, with 19 refractory patients (25%) —defined as lack of any response and/or progression within the first 6-month (**Figure 1A**). In the TCE cohort, the median PFS was 5.3 months, with 50% of refractory patients (**Figure 1B**). Consistent with prior reports,^26,27^ the presence of EMD at the time of the anti-BCMA therapy and prior exposure to anti-BCMA therapy was associated with shorter PFS in the CART cohort, but not in the TCE (**Supplementary Figure 1A-B; Supplementary Table 3**). High (>100 ng/mL) soluble BCMA (sBCMA) was associated with poor outcomes in the CART cohort, where this feature was assessed (p=0.042) (**Supplementary Figure 1C**).^26^ Using the MyCARe score, available for 58 (78%) patients treated with CART, the high-risk group identified a very small population of RRMM with primary refractoriness to anti-BCMA CART (n=3; p<0.0001; **Supplementary Figure 1D**).^22^ However, no differences were observed between those identified as low and intermediate risk groups (p=0.1). Cilta-cel showed a trend toward better PFS compared to ide-cel, that did not reach statistical significance (**Supplementary Figure 1E**). No difference between anti-BCMA antibodies was observed (**Supplementary Figure 1F**).

**Figure 1.**
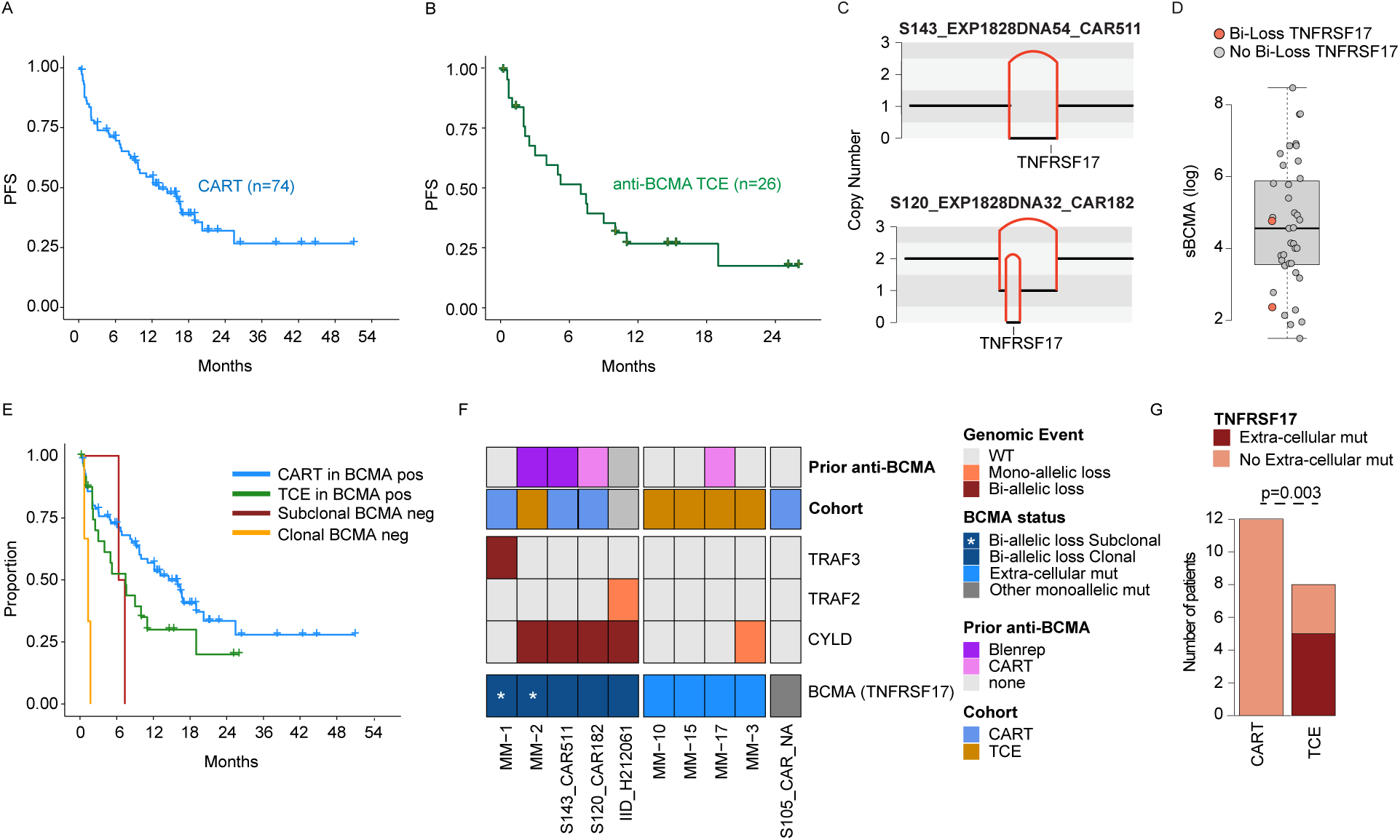
Study population clinical outcomes. **A-B)** Kaplan-Meier showing progression-free survival (PFS) for the 74 and 26 patients with samples collected before CART (**A**) and TCE (**B**). **C)** Copy number and structural variants plots for the two patients with biallelic deletions on *TNFRSF17* before CART/TCE treatment. The horizontal black lines represent the total copy number, the vertical red lines the SV deletion breakpoints. **D)** The impact of biallelic deletions on soluble BCMA (sBCMA). **E)** Kaplan-Meier showing the impact of clonal and subclonal *TNFRSF17* biallelic loss before treatment with CART and TCE in terms of PFS. The p-value was generated using the log-rank test. **G)** Prevalence of mutations impacting anti-BCMA therapy binding among post-CART and post-TCE samples.

### Loss of TNFRSF17 (BCMA) after anti-BCMA therapy

Combining the CART and TCE series, we observed 6 patients with biallelic loss of *TNFRSF17* (**Figure 1C**). Of these, 4 had detectable biallelic loss of *TNFRSF17* before anti-BCMA CART, and two of them had a complete clonal loss of *TNFRSF17* and were previously exposed to anti-BCMA therapies (Belantamab mafodotin and anti-BCMA CART). Interestingly, one of these two patients had low sBCMA, while the other patient had measurable sBCMA levels close to the group median (117 ng/mL; **Figure 1D**). This discordance may be explained by the significant tumor burden (Metabolic Tumor Volume^26^ >240; beta 2 microglobulin= 9) in this second patient, and points to the presence of a residual reservoir of tumor cells with retained *TNFRSF17*. This example suggests that the use of sBCMA to predict loss of *TNFRSF17* may be insufficient in isolation and needs to be integrated with other factors, including radiologically assessed tumor burden, as we have previously shown.^26^ Subclonal BCMA losses were detected using single-cell WGS in one patient previously exposed to Belantamab, and using bulk WGS in another not exposed to any anti-BCMA therapy.^13^ As expected, patients with clonal biallelic loss of *TNFRSF17* before anti-BCMA CART/TCE treatment were characterized by primary refractoriness and poor outcomes. The two other patients with subclonal biallelic loss progressed within 7 months and relapsed disease was associated with the selection of the BCMA-negative clone detected at baseline (**Figure 1E**). Two additional patients with *TNFRSF17* diploid status at baseline acquired biallelic loss of *TNFRSF17* at relapse following anti-BCMA CART or TCE therapy. Overall, these data support the idea that biallelic loss of BCMA occurs in a relatively small fraction of MM, and the risk of this event can be increased by prior exposure to other anti-BCMA therapies—especially agents associated with chronic exposure (e.g. Belantamab). Interestingly, all patients harboring biallelic loss of BCMA also carried biallelic inactivation of *CYLD* or *TRAF3*, suggesting a compensatory mechanism that maintains NF-κB pathway activation, typically supported by BCMA signaling (**Figure 1F**).

### Tumor evolution after CART and TCE treatment

WGS data from samples collected at the time of progression after CART and TCE therapy were analyzed in 12 and 8 patients, respectively. Among the post-CART samples, we identified only one nonsynonymous mutation in the extracellular domain of *TNFRSF17*, specifically the P33S mutation in post-CART samples from a patient treated with orva-cel (55289_CG22_1561). Additionally, this patient had a whole chromosome 16 monoallelic loss. The P33S cancer cell fraction was 20%, reflecting a subclonal status. Prior functional studies in human cell lines demonstrated that this mutation had a neutral effect on anti-BCMA TCEs and ide-cel MM binding and killing activity.^13^ To investigate its impact in the context of orva-cel and cilta-cel treatment we similarly employed different human cell lines and confirmed that the P33S mutation does not affect CART binding of BCMA (e.g.,**Supplementary Figure 2**). The absence of mutations affecting CART BCMA binding at relapse is consistent with recent findings in a cohort of 24 post-CART patients, where circulating free DNA analyzed via targeted sequencing similarly showed no evidence of such mutations.^16^ In striking contrast to the post-CART setting, 5 out of 8 patients in the TCE group exhibited clonal mutations in the BCMA extracellular domain or TCE binding site, and one patient showed biallelic loss of BCMA (compared to 0/12 in the CART group; p=0.003, Fisher’s exact test; **Figure 1G**).

Nine of the 12 patients with available post-CART samples also had a sample collected immediately before CART therapy, and three had a third sample collected at day +30. Using WGS data from these serially collected samples, we reconstructed phylogenetic trees to examine patterns of selection and genomic evolution under CART selective pressure. Despite the short PFS between sequential samples (8/9 progressed within the first 6 months), 5 of 10 patients exhibited clonal changes upon progression, while the other 50% maintained a relatively stable clonal structure (**Supplementary Figure 3**). In all patients who experienced branching evolution at progression, the dominant clone driving progression was always detectable in the pre-CART sample. This contrasts with TCE, where BCMA-mutated clones are usually undetectable until approximately six months of therapy.^15^ In the three patients with samples collected on day +30 post-CART infusion, no significant clonal changes were observed. Notably, and in contrast to TCE, no recurrent genomic events were found in the dominant clones selected at relapse among the 12 post-CART samples.

### Genomic complexity and high proliferation drive plasma cell escape, promoting resistance to anti-BCMA CART cells and TCE

Considering the relatively low prevalence of genomic events affecting *TNFRSF17* and the relative lack of recurrent genomic drivers selected at progression in patients with suboptimal response and/or primary refractory disease, we hypothesized that resistance to CART might be driven by pre-existing genomic drivers. To comprehensively investigate these important aspects, we leveraged the WGS from 74 and 26 patients treated with CART and TCE, respectively, with samples available before therapy. After annotating each sample according to the recent myeloma genomic classification,^28^ we observed a significant difference in PFS between patients treated with CART harboring a complex vs simple genomic profile, where complex is defined as the presence of 6 or more genomic events involving tumor suppressor genes or high risk features (p=0.033; **Figure 2A**). All the TCE but one patient were defined as complex, and the only case with a simple profile was still in remission after 300 days. As the association between genomic complexity and resistance to CART has also been reported in large B-cell lymphomas (LBCL),^29,30^ we interrogated the role of genomic complexity in promoting resistance to CART therapy using a validation set of 35 patients treated with cilta-cel who had MSK-IMPACT targeted sequencing data available (**Supplementary Table 4**).^31^ Despite the limited sample set and targeted panel limited genomic resolution, we still observed a strong association between genomic complexity and PFS (p=0.0016; **Figure 2B**). Collectively, these data showed that, similarly to LBCL,^29,30^ genomic complexity is a critical marker of decreased sensitivity to anti-BCMA CART in RRMM.

**Figure 2.**
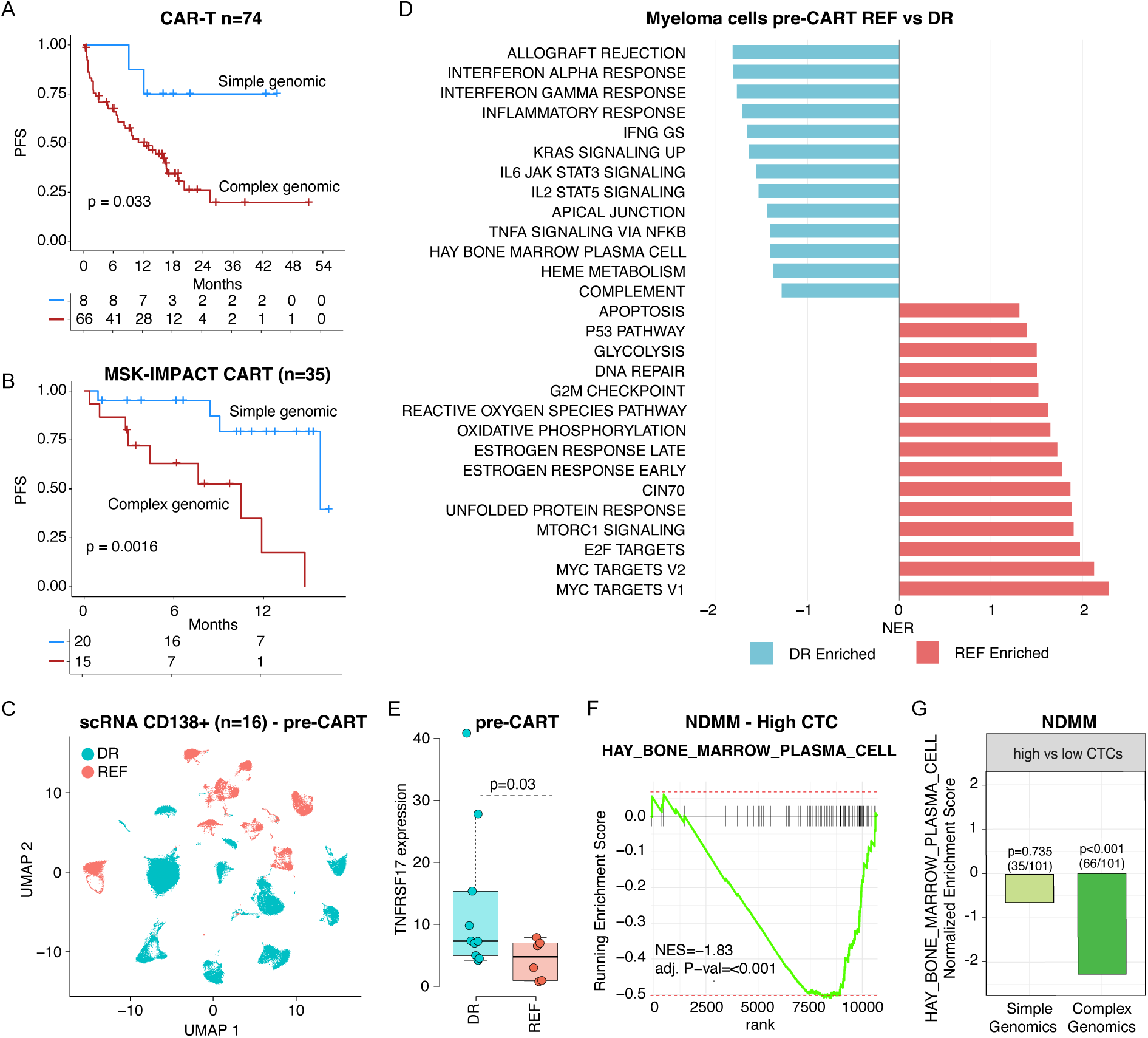
Genomic features associated with poor outcomes after anti-BCMA CART therapy. **A)** Kaplan-Meier analysis showing progression-free survival (PFS) according to genomic complexity (as defined by Maura et al., *JCO* 2024) prior to anti-BCMA CART therapy. **B)** Kaplan-Meier analysis of PFS in the MSK-IMPACT-Hem validation cohort stratified by genomic complexity before CART therapy. **C)** UMAP representation of single-cell multiple myeloma profiles from 16 patients. **D)** Gene set enrichment analysis (GSEA) comparing refractory (REF) versus durable responders (DR) after CART therapy. **E)** Boxplot depicting *TNFRSF17* expression differences between responders and refractory patients. **F)** GSEA of plasma cell signatures in newly diagnosed multiple myeloma (NDMM) patients with high circulating tumor cells (CTCs). **G)** Enrichment analysis showing loss of plasma cell signatures in patients with both high CTCs and high genomic.

### Tumor and immune transcriptomic signatures associated with TCE failure

To further investigate the link between anti-BCMA therapy and genomically complex MM, we analyzed single-cell RNA sequencing (scRNAseq) data from clonal plasma cells in 16 patients collected prior to CART therapy (**Fig 2C**). Of these, 5 also had matched WGS data (2 and 3 with high and low genomic complexity, respectively).

Refractory patients (n=6)^32^ showed a significant increase in proliferation, genomic instability and downregulation of plasma cell signatures compared to durable responders (n=10, **Figure 2D; Supplementary Table 5-6**). In keeping with this phenotype and in line with prior observations,^32^ *TNFRSF17* expression was also reduced in the refractory group (p=0.03; **Figure 2E**). The association between high genomic complexity, increased proliferation, and reduction of the plasma cell transcriptional program is defined here as “plasma cell identity escape”. The idea that genomically unstable and highly proliferative MM cells diverge from the plasma cell transcriptional and biological background has been observed with other targeted therapies in MM.^33–35^ This is also supported by clinical observations in MM patients who progressively reduce protein production while increasing proliferation and the ability to form tumor masses. As anti-BCMA therapies are rapidly moving into frontline treatment, we sought to investigate whether these patterns of plasma cell identity escape can already be detected in newly diagnosed MM (NDMM). By analyzing 540 NDMM patients with available RNAseq, WGS and WES,^36^ we observed a strong association between high proliferation, elevated circulating tumor cells (CTC), genomic complexity, and downregulation of the plasma cell transcriptional program (**Fig. 2F-G; Supplementary Table 7**). Overall, these data define a population of MM in which the loss of plasma cell identity promotes primary resistance to therapies that depend on plasma cell biology, including immunomodulatory drugs (IMiDs), anti-BCMA, and anti-CD38 agents.^37^

### Immune and genomic profiles are interlinked in promoting refractoriness to CART

Distinct patterns of T-cell exhaustion and immune dysfunction have been associated with suboptimal response and early progression after both CART and TCE.^38,39^ To further investigate this, we analyzed immune profiles using scRNA-seq from the 11 patients described above and integrated these data with 6 previously published patients^32^ treated with anti-BCMA CART (**Fig 3A**). Consistent with prior observations, suboptimal response to CART was associated with higher level of activation/exhaustion in the T CD8 cells (**Fig 3B-C; Supplementary Table 8**). The same patterns were not observed in the T CD4 cells (**Supplementary Figure 4**). In line with prior evidence,^33,40–43^ these data suggest a strong interconnection between tumor biology and the immune microenvironment, in which highly proliferative and genomically complex tumors drive rapid expansion and subsequent exhaustion of the surrounding bone marrow immune compartment.

**Figure 3.**
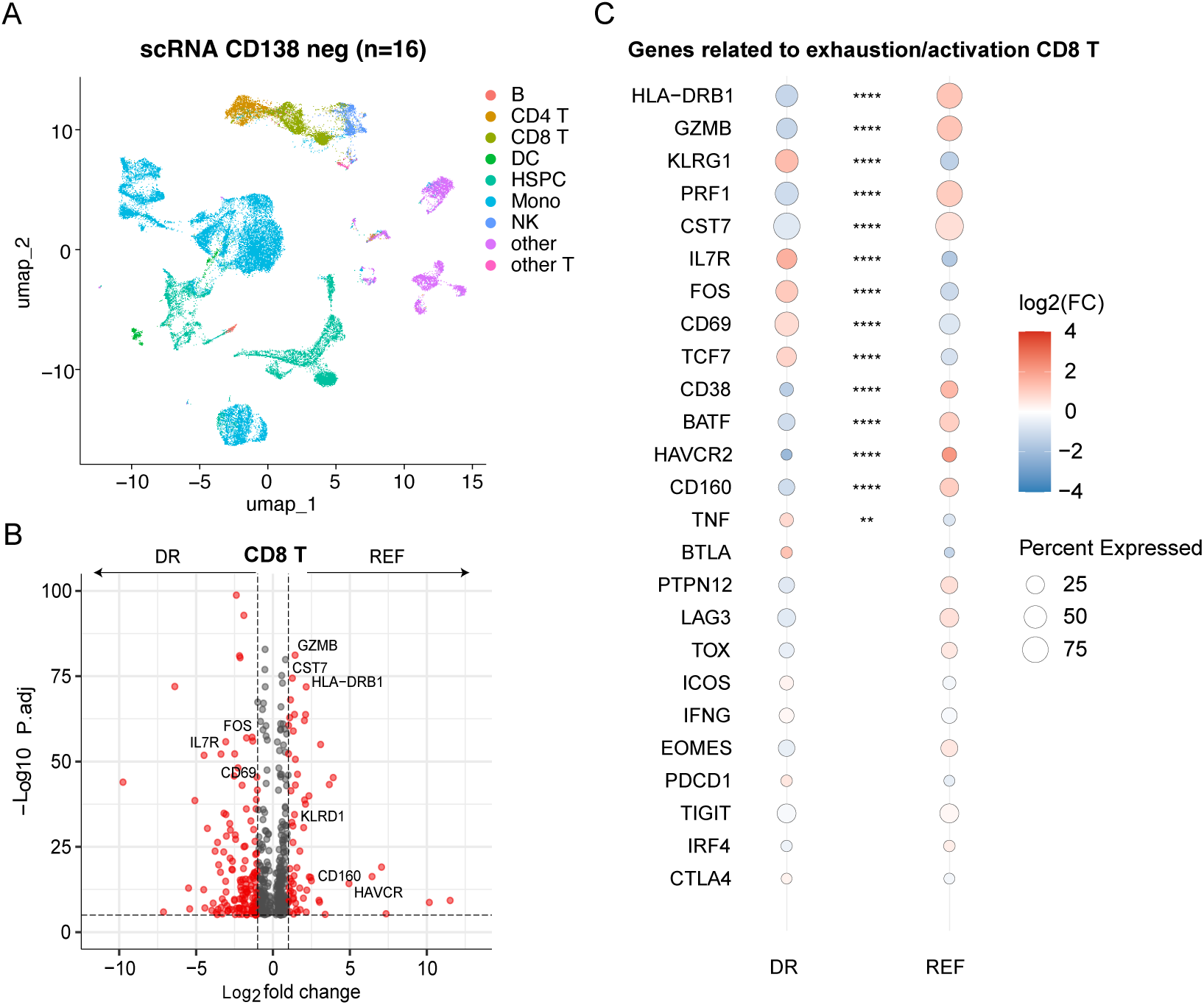
T CD8 exhaustion and activation is associated with refractoriness to anti-BCMA CART. **A)** UMAP summarizing the main population in the CD138-negative cells. A total of 42,957 cells was analyzed**. B-C)** Increased immune exhaustion/activation in the T CD8 of refractory (REF) patients compared to durable responders (DR).

### Loss of genes involved in plasma cell biology associated with refractoriness to anti-BCMA CART and TCE

As genomic complexity and loss of plasma cell identity are associated with refractory disease and poor response after CART and TCE, and likely promote high tumor burden and immune dysfunction, we went on to investigate the clinical impact of established MM and cancer driver genes, identifying two main groups (**Figure 4A; Supplementary Table 9**). Consistent with our phenotypic observations, the first group included loss of function genes involved in plasma cell identity and differentiation. In addition to biallelic loss of *TNFRSF17*, genomic loss of the following genes involved in B-cell and plasma cell biology was associated with shorter PFS and refractoriness: *XBP1*, *CD38*, *IKZF3*, and *POU2AF1*. Interestingly, 14 out of 17 patients with EMD harbored at least one of these lesions. Loss of *CD38* has been reported to potentially confer resistance to anti-CD38 monoclonal antibodies through reduced *CD38* gene and protein expression. In line with recent findings,^35,44^ 24% of patients exhibited *CD38* loss, which was associated with poor outcomes after anti-BCMA CART therapy (p=0.0024; **Figure 4B-C; Supplementary Figure 5A-B**). To better characterize the link between the loss of function of *CD38* and *TNFRSF17* expression, we analyzed gene expression data by integrating 767 NDMM patients enrolled in the CoMMpass trial with an independent Moffitt cohort comprising 269 NDMM and 683 RRMM.^45^ A strong direct correlation was observed between *TNFRSF17* and *CD38* in both the RRMM and NDMM (p<0.0001; **Supplementary Figure 5C-D**). The presence of focal loss of *IKZF3* was associated with primary refractoriness to both CART and TCE therapies (p=0.0014; **Figure 4E**). This deletion has recently been linked to resistance to anti-CD38 antibodies and IMiDs, and although it has a limited impact on *TNFRSF17* expression in NDMM, it emerges as a potential high-risk feature in both NDMM and RRMM treated with immunotherapy. Loss of *XBP1* was also associated with poor outcomes after anti-BCMA CART, showing a strong correlation with *TNFRSF17* in gene expression (**Supplementary Figure 5E-G**). Finally, we identified a patient who was refractory to anti-BCMA TCE therapy despite not exhibiting elevated levels of sBCMA. This patient harbored a unique breakage-fusion-bridge event involving *TNFRSF13B* (i.e., *TACI*; **Figure 4F**). Notably, the same patient achieved a prolonged remission following treatment with an anti-GPRC5D TCE, suggesting primary refractoriness to anti-BCMA immunotherapy, but not to immunotherapy in general. Overall, patients harboring any of these events affecting plasma cell identity experienced shorter PFS compared with the others (p<0.0001; **Figure 4G; Supplementary Figure 5H-I**).

**Figure 4.**
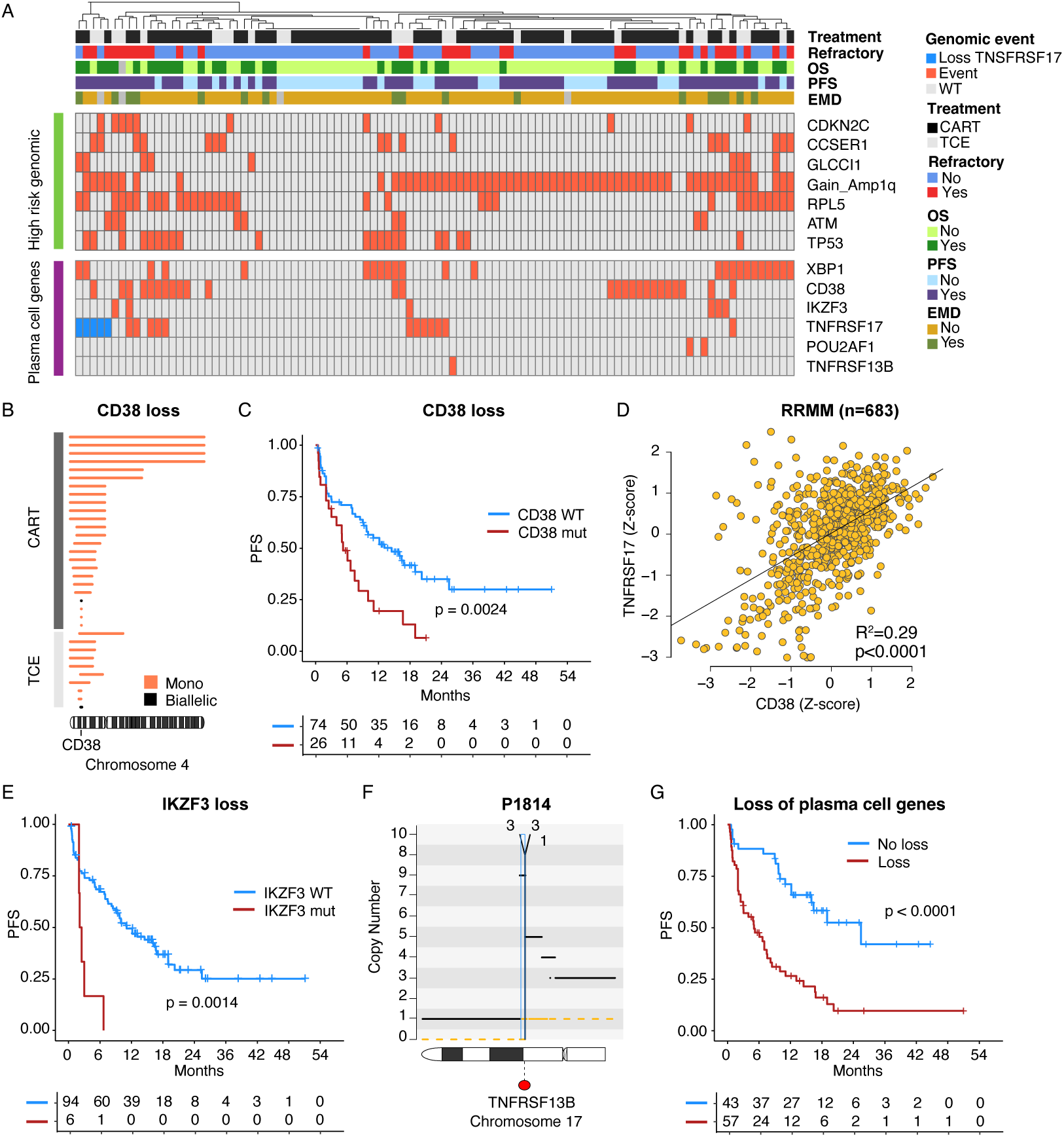
Loss of genes involved in plasma cell biology associates with refractoriness to anti-BCMA CART/TCE. **A)** Heatmap summarizing distribution, prevalence and patterns of co-occurrence of all genomic features with significant impact on PFS post-anti-BCMA CART. **B)** Summary of all the deletions involving *CD38* across our cohort. Each segment represents a patient and the length the size of the deletion. **C)** Kaplan-Meier showing the impact of *CD38* loss before treatment with anti-BCMA CART/TCE in terms of progression-free survival (PFS). **D)** Correlation between *CD38* and *TNFSRF17* expression across 683 RRMM with available RNAseq, respectively. **E)** Kaplan-Meier showing the impact of *IKZF3* loss before treatment with anti-BCMA CART in terms of PFS. **F)** Copy number and structural variants (SV) plots for one patient with chromothripsis causing high copy number gains in *MAP3K14*. The horizontal black and orange lines represent the total copy number and the minor allele. The vertical red, green, blue and black lines represent the following SVs: deletions, duplications, inversions and translocations. **G)** Kaplan-Meier showing the impact of loss of genes involved in plasma cell biology before treatment with anti-BCMA CART in terms of PFS

### High risk genomics and TP53 promote primary refractoriness to anti-BCMA CART and TCE

The second predictive group included events known to confer high risk disease such as gain1q, mutations in *TP53*, deletions of *CDKN2C* (1p32.3) and *RPL5* (1p22), loss of *CCSER1, ATM,* and *MAF*/*MAFB* translocations, and strongly correlate with shorter PFS and refractory disease compared to wild-type (**Figure 4A**; **Figure 5A-B; Supplementary Figure 6A-D**). Importantly, 15/17 patients with EMD overlapped with this group of patients with high risk genomic profile. Among these features, the presence of *TP53* mutation showed the most striking association in both CART and TCE, even when the two datasets were considered separately (**Supplementary Figure 6C-D**). To better characterize the impact of this recurrent alteration, we investigated the association between *TP53* and *TNFRSF17* at the transcriptional level. To do so, we examined bulk RNA-seq correlations in both NDMM and RRMM.^45^ In both groups, *TP53* expression was significantly correlated with *TNFRSF17* (p<0.0001; **Figure 5C; Supplementary Figure 6E**). Overall, patients with high-risk genomic features experienced significantly poorer outcomes compared with those without these alterations (p=0.0002; **Figure 5D**). Importantly, the presence of high-risk genomic features and loss of plasma cell identity genes were both independently associated with inferior outcomes in a multivariate model that included EMD, age, prior exposure to anti-BCMA therapy, number of therapy lines, and product type (**Figure 5E**).

**Figure 5.**
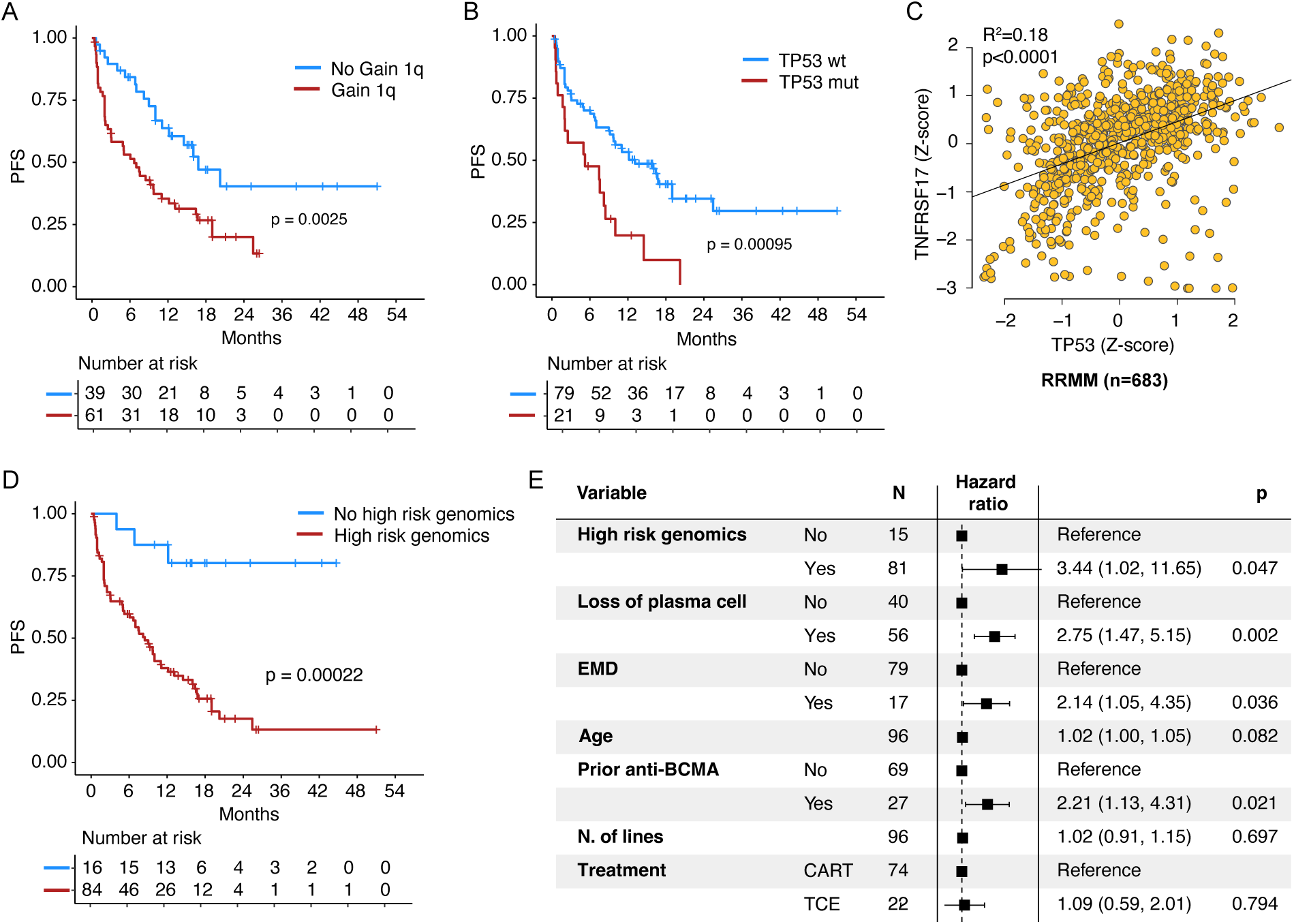
*TP53* promotes resistance to anti-BCMA CART therapy. A-B) Kaplan-Meier analyses showing the impact of gain 1q (**A**) and nonsynonymous *TP53* mutations (**B**) prior to anti-BCMA CART therapy on progression-free survival (PFS). **C)** Correlation between *TP53* and *TNFRSF17* expression across 683 RRMM samples with available RNA-seq data. **D)** Kaplan-Meier analysis showing the impact of loss of genes involved in plasma cell biology on PFS following anti-BCMA CART therapy. **E)** Multivariate Cox regression model assessing the impact of plasma cell gene loss and high-risk genomic features, adjusted for extramedullary disease (EMD), age, prior anti-BCMA therapies, number of prior lines, and treatment product (CART vs TCE).

### Loss of TP53 confers suboptimal response to anti-BCMA CART therapy

To further investigate the association between loss of plasma cell identity, genomic complexity and suboptimal response to anti-BCMA redirecting therapies, we leveraged recently developed Bclγ1, MIcγ1, and TP53–Bclγ1 mouse models of MM. ^46,47^ The first two models were *Trp53* wild-type, whereas the latter was characterized by monoallelic loss of *Trp53* (**Figure 6A–B**; see **Methods**). Consistent with the human data, scRNAseq analysis revealed that mice with *Trp53* loss exhibited a significant downregulation of *Tnfrsf17* compared to the wild-type models (**Figure 6C; Supplementary Table 6**). Mirroring the results observed in MM patients, MM cells from Bclγ1 and MIcγ1 mice were also more enriched for plasma cell gene-expression signatures compared with TP53–Bclγ1 mice (**Figure 6D**). Overall, these experimental findings provide experimental support that *TP53*-deficient MM exhibits downregulation of *Tnfrsf17*, likely reflecting both enhanced genomic instability and loss of plasma cell identity.

**Figure 6.**
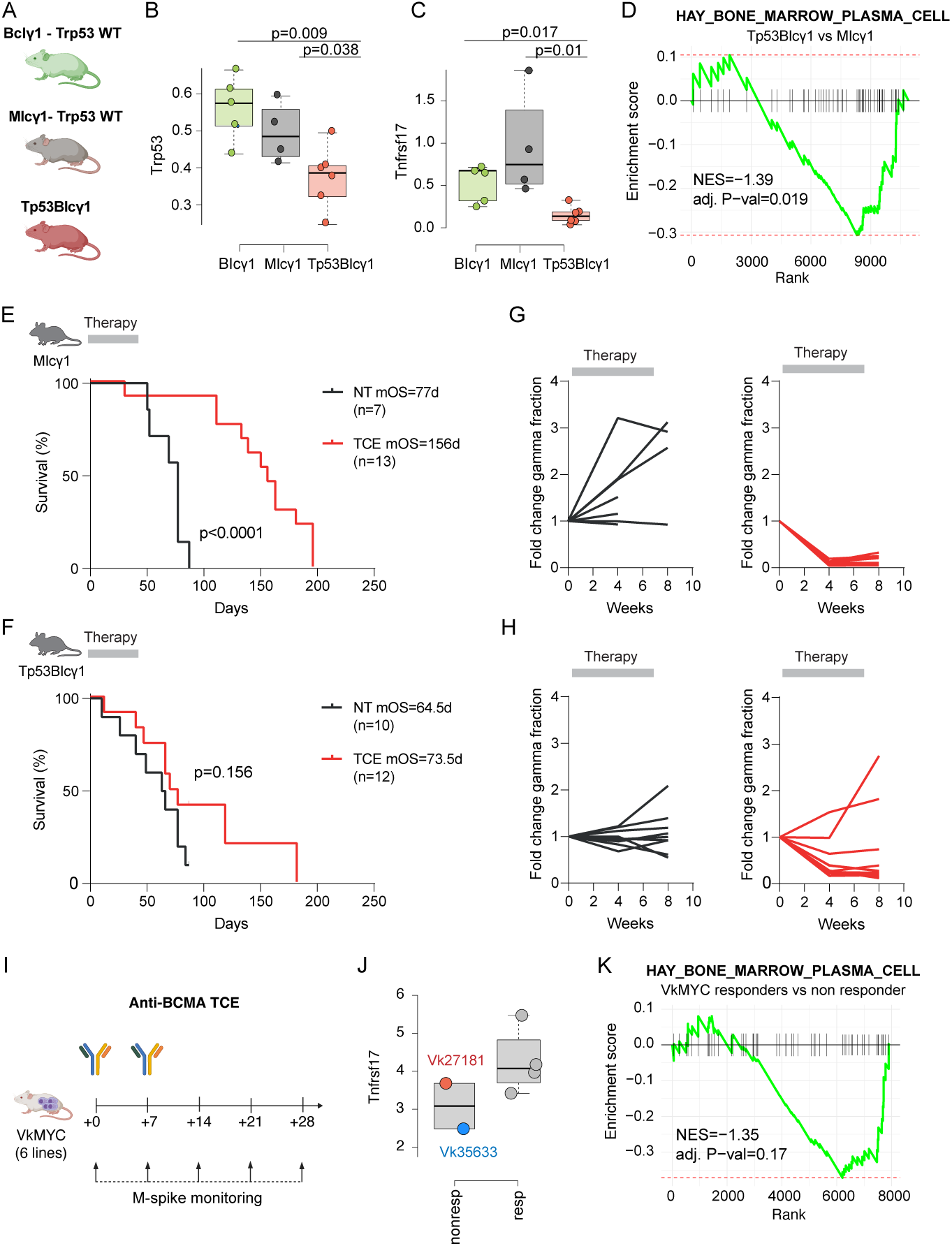
*Trp53* loss and reduced plasma cell gene expression signatures impair response to anti-BCMA TCE in multiple myeloma mouse models. A) Schematic of the Bclγ1, MIcγ1, and Tp53BIcγ1 mouse study design. **B-C)** Differential expression of *Trp53* (**B**) and *Tnfrsf17* (**C**) across the three mouse models. Gene expression was derived from pseudo-bulked scRNAseq data. **D)** Enrichment of plasma cell gene expression signatures in MIcγ1 compared to the Tp53BIcγ1 mice. **E–H)** Response of Tp53BIcγ1 and MIcγ1 mice to anti-BCMA TCE treatment. **I)** Study design showing treatment of six distinct VkMYC transplantable lines with anti-BCMA TCE. **J)** The two refractory VkMYC lines exhibited downregulation of *Tnfrsf17* compared to responders. **K**) The same refractory VkMYC lines showed significant downregulation of the plasma cell gene expression signature.

To test responses to anti-BCMA therapy in vivo, a murine BcmaxCd3 surrogate TCE antibody was administered to two MM mouse strains with wild-type and deleted *Trp53* gene (i.e., MIcγ1 and Tp53-BIcγ1 mice, respectively). Therapy was administered for 8 weeks once MM was observed. MIcγ1 mice with wild-type *Trp53* exhibited complete response to the TCE at week 4 post-treatment initiation in comparison to isotype-treated mice, which was sustained until week 8 in all mice (**Figure 6E-F**). In contrast, responses to TCE in Trp53-BIcγ1 mice with heterozygous deletion of *Trp53* were not as deep and prolonged, and 2 of 12 mice relapsed during treatment (**Figure 6G-H** and **Supplementary Figure 7A**). Accordingly, MIcγ1 mice receiving BcmaxCd3 therapy showed a major extension in survival with respect to control animals (median overall survival [OS], 156 vs 77; p<0.0001), while Tp53-BIcγ1 mice only showed a modest increase in survival with respect to controls (median OS, 73.5 vs 64.5; p=0.068). These results experimentally validate the data in MM patients, reinforcing the concept that the *TP53* status impacts responses to BCMA-targeted agents.

To further validate the association between the loss of plasma cell identity and resistance to anti-BCMA immunotherapy, we employed a second mouse model. We investigated six independent Vk*MYC mouse MM lines treated with an anti-BCMA TCE (**Figure 6I; Methods**).^48–50^ Two of the six Vk*MYC lines exhibited primary refractoriness to anti-BCMA TCE treatment in vivo (**Supplementary Figure 7B-C**). Analysis of RNA-seq data from the two refractory lines with available data, particularly Vk35633, demonstrated low *Tnfrsf17* expression (**Figure 6J**) as well as a marked reduction in the plasma cell gene expression signature (**Figure 6K**). Overall, these findings in the Vk*MYC mouse support a model in which aggressive and genomically unstable MM are less sensitive to anti-BCMA TCE therapy due to downregulation of their plasma cell identity, including BCMA expression, thereby evading treatment pressure.

## DISCUSSION

Although anti-BCMA CART-cell therapies have shown unprecedented efficacy in MM, most patients eventually relapse, and the potential for truly durable remissions remains unclear.^3,4,7,21,22,51–53^ Several studies have demonstrated that this clinical behavior is associated with distinct patterns of immune exhaustion in the tumor microenvironment, in particular in the T CD8.^38,39,54^ These exhaustion/activation programs have been consistently linked to poor outcomes in MM, even with other immunotherapies such as anti-CD38 monoclonal antibodies.^33,43^ In parallel to these discoveries, recent investigations have highlighted the critical role of genomic complexity and instability in promoting resistance to MM therapies.^28,33,42,43,55^ Given that tumor cells and immune components co-exist within a highly interactive ecosystem, it is likely that genomic aggressiveness—defined by high proliferation and instability—and immune exhaustion are interrelated phenomena. Consistent with these observations, our study identified a significant association between genomic instability, proliferation, T-cell exhaustion, EMD, and poor outcomes following either CART-cell or TCE therapy. By integrating these findings, we propose the existence of a distinct evolutionary trajectory in MM toward a highly proliferative and genomically unstable state that progressively loses its dependence on plasma cell identity. This process may be driven by therapeutic selection pressures (e.g., loss of *CD38* after daratumumab exposure^56^) or may already exist at baseline in high-risk patients (e.g., high circulating tumor cells with genomic complexity).^36^ Because most current MM therapies target plasma cell biology, these transformed subclones become increasingly resistant and less sensitive to such approaches (**Figure 7**). Consequently, these patients often exhibit primary refractoriness to T-cell-redirecting therapies targeting BCMA, which is frequently downregulated together with other plasma cell-specific genes. This phenomenon is not entirely new. Clinically, MM has long been recognized to evolve from a bone marrow-restricted, immunoglobulin-producing disease to an aggressive, extramedullary, or leukemic phase characterized by rapid proliferation and minimal immunoglobulin secretion.^57,58^ Based on our genomic findings and prior historical clinical observations, we propose the term “plasma cell identity escape” to describe this state, defined by the loss of plasma cell features and associated therapeutic vulnerabilities. While larger patient cohorts are needed to accurately refine the definition of this entity, our findings mark a conceptual step forward for the MM field. Importantly, nearly all emerging MM therapies are plasma cell-oriented and will likely be ineffective in this subgroup, as exemplified by anti-BCMA CART and TCE resistance. Future research should therefore redefine therapeutic paradigms and explore alternative strategies that target MM cells independently of plasma cell lineage features. This shift will be crucial not only for relapsed/refractory MM but also for newly diagnosed disease, as our data show that plasma cell identity escape can already be detected in some NDMM cases and is associated with early resistance to IMiDs and anti-CD38 therapies.

**Figure 7.**
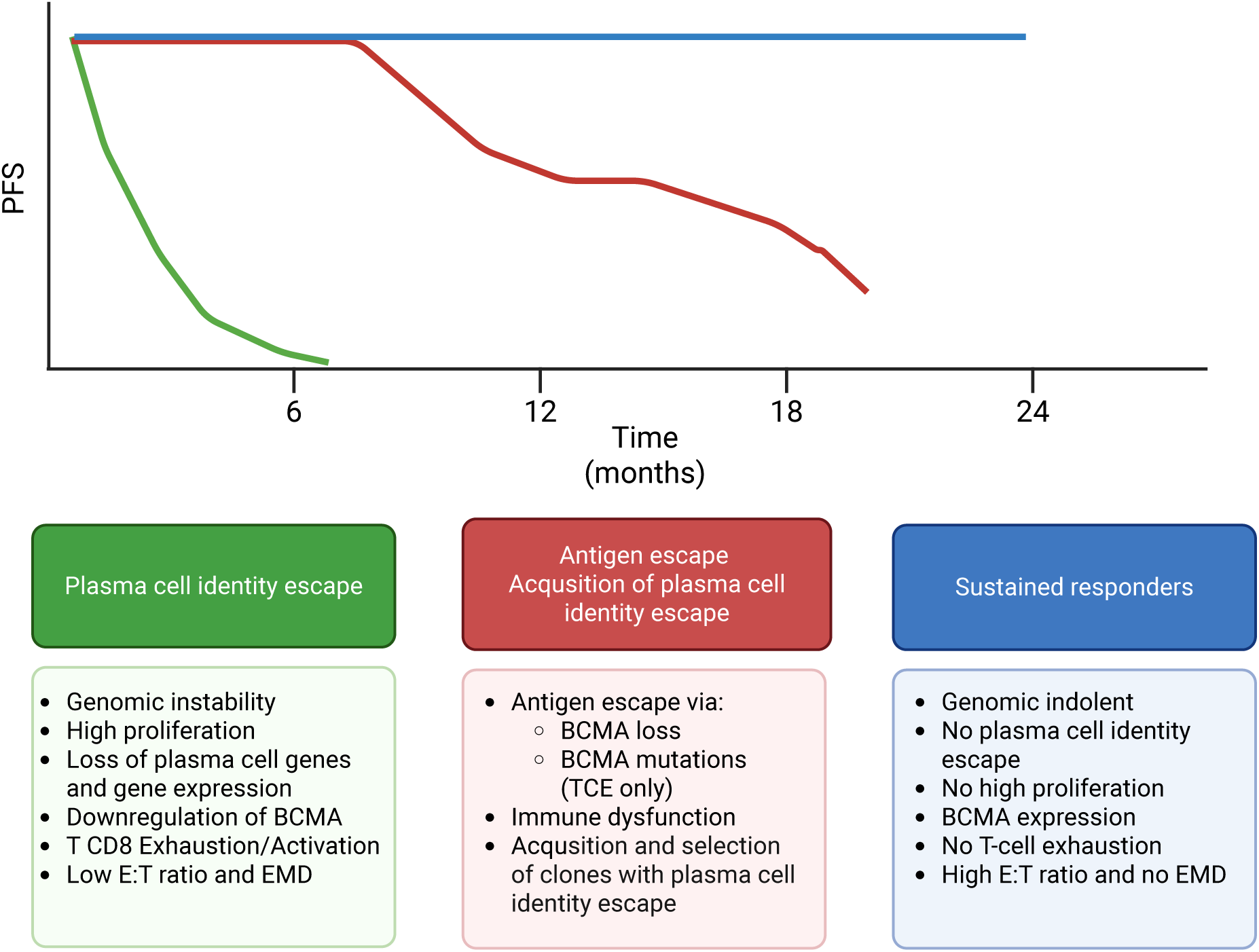
Overview of patient subgroups showing distinct clinical outcomes and corresponding genomic alterations after anti-BCMA therapies.

The association between plasma cell identity escape, EMD, and CD8⁺ T-cell exhaustion or activation—two of the most established prognostic factors in anti-BCMA CART-cell and TCE therapies—aligns with the model described above. Aggressive and highly proliferative tumors exert greater pressure on the immune system than indolent ones, contributing to increased tumor burden and facilitating extramedullary growth. This observation is critical because, although the interplay between immune and tumor compartments requires further functional validation, our data suggest a predominantly tumor-centric mechanism.^42^ Specifically, the occurrence of antigen escape in most patients who initially respond but later progress,^13^ together with the loss of plasma cell identity in those with primary refractoriness, supports a model in which the tumor itself drives resistance, thereby promoting immune exhaustion, EMD development, and uncontrolled proliferation. Within this framework, strategies aimed solely at reinvigorating T cells in vitro may prove insufficient unless accompanied by effective tumor control and suppression of proliferative dynamics.

Comparing WGS data from patients who relapsed after anti-BCMA CART and TCE therapies, we observed a striking absence of *TNFRSF17* mutations in the BCMA extracellular domain in the post-CART setting, in contrast to their frequent detection after anti-BCMA TCE in our study (5/8 cases) and in others recently presented. Prior studies indicate that such mutations arise under prolonged selective pressure from continuous TCE exposure.^13–15^ This discrepancy likely reflects fundamental differences in therapeutic kinetics: CART cells exert a strong, short-term selective pressure, whereas TCE therapies apply sustained pressure that promotes antigen escape (**Figure 7**). This distinction also challenges the perceived role of CART persistence in driving relapse dynamics. If long-term CART activity were a dominant selective force, one would expect to observe similar antigen escape mutations as seen with TCE therapy. The lack of such mutations suggests that persistent CART pressure is not the primary mechanism of relapse in this context.

Overall, our study provides evidence that integrated genomic and transcriptomic profiling can identify a subset of MM patients with plasma cell identity escape, characterized by genomic instability, proliferative drive, and resistance to therapies targeting plasma cell biology. Recognizing and defining this entity is essential for developing new therapeutic strategies capable of overcoming resistance in this aggressive form of myeloma.

## METHODS

### Patients

The Moffitt samples were obtained with approval from the Moffitt Institutional Review Board and an external IRB (Advarra). All patients provided written informed consent under one of two IRB-approved protocols: the Total Cancer Care (TCC) institutional biorepository protocol (MCC#14690; Advarra IRB Pro00014441) or a dedicated biospecimen collection protocol (Pro00021733). Consented patients agreed to the collection of additional bone marrow aspirates during clinically indicated biopsies, peripheral blood samples, and access to their medical records. This study, adhering to the Declaration of Helsinki, received approval from all institutions which collected samples and data for this study: Moffitt Cancer Center, Calgary University and Memorial Sloan Kettering Cancer Center. All samples and clinical data were collected with patients’ consent.

### WGS and WES sequencing and data analysis

Bone marrow samples were collected from 89 patients treated with either anti-BCMA CART (n=75) and/or anti-BCMA TCE (n=15). All samples were sorted using the StemCell EasySep Human CD138 positive selection kit II (StemCell, Cambridge, MA). The positive CD138+ were submitted for either WGS or WES, using peripheral blood mononuclear cells as normal matched. WGS was carried out by the Molecular Genomics Core Facility at Moffitt Cancer Center contingent on the availability of high-quality pre-treatment samples. Germline DNA was obtained from peripheral blood mononuclear cells, while tumor DNA was extracted from CD138-enriched bone marrow aspirate specimens using magnetic bead-based selection. Genomic DNA was isolated using the AllPrep DNA/RNA/miRNA Universal Kit (Qiagen). For library preparation, 100 ng of genomic DNA was fragmented using the Covaris ML230 focused ultrasonicator (Covaris, Inc.), followed by library construction with the TruSeq DNA Nano Library Prep Kit (Illumina) in accordance with manufacturer instructions. Library quality and fragment size distribution were assessed via Agilent TapeStation, and quantification was performed using the Kapa Library Quantification Kit (Roche). Sequencing was conducted on the Illumina NovaSeq 6000 system using S4-300 reagent kits with a 2×150 bp paired-end configuration. Tumor samples were sequenced to a median coverage of 60×, and matched germline controls to 30×.

For the MSKCC samples, DNA was extracted from bone marrow following CD138+ magnetic bead selection. Matched normal samples were collected from peripheral blood mononuclear cells. For each WGS sample, 500ng of genomic DNA was sheared, and sequencing libraries were prepared using the KAPA Hyper Prep Kit (KAPA Biosystems). Samples were run on an Illumina NovaSeq 6000 in a 150bp/150bp paired-end runs, using the SBS v1 Kit and an S4 flow cell (Illumina).^59^

For the Calgary samples, WGS was performed as previously reported at the New York Genome Center. ^13,14^

All FASTQ were aligned to the GRCh38.p14 genome reference. WGS data from Moffitt were processed using DRAGEN Somatic v4.2.4 for alignment and mutation calling. Structural variants were identified using Manta ^60^, Svaba ^61^, IgCaller ^62^, and Delly2 ^63^ as part of the mgp1000 somatic pipeline (v2.1, commit 239cf3c). MSKCC and Calgary samples were processed using the mgp1000 preprocessing and somatic pipeline (v2.1, commit 239cf3c; https://github.com/UM-Myeloma-Genomics/MPG). All samples underwent CNV calling with ASCAT v3.1.1 using alleleCounter v4.3.0.^64^ Clonality analysis was performed using dpclust3p v1.0.8 and DPClust v2.2.8.^65^ Mmsig was used to estimate mutational signatures. The genomic classification and definition of “complex status” are available on GitHub: https://github.com/UM-Myeloma-Genomics/GCP_MM.

### MSK IMPACT

MSK-IMPACT-HEME (Integrated Mutation Profiling of Actionable Cancer Targets) identifies specific mutations in 468 genes.^66^ DNA was extracted from either from bone marrow following CD138+ magnetic bead selection, or from plasmacytoma biopsy samples. Matched normal controls were collected from nail clippings or saliva. Mutations were detected by hybridization capture of DNA followed by massively parallel sequencing on an Illumina NovaSeq 6000 instrument, while copy number calling uses the FACETS algorithm. De-identified data is publicly available at www.cbioportal.org.

### Soluble BCMA (sBCMA) Quantification

For patients with available serum samples collected prior to lymphodepletion (within day -7 to day -5 prior to initiation of chemotherapy), soluble BCMA (sBCMA) levels were measured using a commercial ELISA kit (R&D Systems, MN, USA; #DY193). Serum samples were serially diluted (between 0-500x) and plates were developed per the manufacturer’s protocol. Absorbance was measured using a Cytation3 imaging plate reader (Biotek) with Gen5 software v2.09. sBCMA concentrations were interpolated from a standard curve using GraphPad Prism v10.

### Bulk RNA sequencing

Patients included in this component of the analysis were enrolled in the Total Cancer Care (TCC) protocol, the institutional biobanking study at Moffitt Cancer Center (MCC#14690; Advarra IRB Pro00014441). All participants provided informed consent for donation of research blood and bone marrow (BM) aspirate samples, along with access to clinical data. A total of 716 patients donated 952 BM samples representing active multiple myeloma (269 newly diagnosed, 349 early relapsed/refractory, and 334 late relapsed/refractory) were collected under this protocol. Samples with ≥1 million viable CD138+ cells were analyzed under the Oncology Research Information Exchange Network (ORIEN) AVATAR initiative. Exclusion occurred only in cases of inadequate cell yield or technical assay failure. Fresh BM aspirates were enriched for CD138+ plasma cells using magnetic-activated cell sorting (MACS) with anti-CD138–conjugated microbeads (Miltenyi Biotec, 130-051-301). A total of 1 × 10⁶ viably frozen CD138+ cells per sample were shipped for RNA extraction and sequencing. Total RNA was isolated using the RNeasy Plus Mini Kit (Qiagen), yielding an average cDNA insert size of 216 base pairs. RNA sequencing was conducted using the Illumina TruSeq RNA Exome workflow, including cDNA synthesis, single-library hybridization, and library construction. Sequencing was performed on an Illumina platform using either 100-bp or 150-bp paired-end reads, targeting a minimum of 100 million total reads (50 million paired reads) per sample. Raw FASTQ files were pre-processed to remove adapter sequences and low-quality bases using a multi-step filtering pipeline, including k-mer–based adapter trimming, quality filtering, masking of low-complexity regions, GC and length filtering, and entropy-based exclusion. Cleaned reads were aligned to the GRCh38/hg38 human reference genome using STAR (RRID: SCR_004463), with annotations from GENCODE v32 (RRID: SCR_014966). Gene and transcript-level expression estimates were generated using STAR’s transcriptome alignment output. Expression metrics included estimated read counts, fragments per kilobase of transcript per million mapped reads (FPKM), and transcripts per million (TPM). The expression dataset contained 59,368 entries, of which 19,933 corresponded to protein-coding genes. Only protein-coding genes were retained for further analysis. To prepare data for differential expression and correlation analysis, expression values were transformed using log₂(TPM + 1).

Additional RNAseq data were downloaded from the CoMMpass data set, a data set generated by the Multiple Myeloma Research Foundation: https://research.themmrf.org/. In this data set, CTC counts were estimated using the CellSearch System as previously described.^36^

### Single-Cell RNA Sequencing and Analysis (Baseline Plasma Cell Compartment)

Freshly isolated bone marrow aspirates were processed within hours of collection at the baseline pre-treatment time point (prior to CART infusion). CD138+ and CD138–mononuclear cell fractions were enriched via magnetic bead-based separation. CD138+ cells were encapsulated separately using the Chromium Controller platform (10x Genomics), with 5000 cells targeted per fraction for molecular barcoding and library construction. Libraries were generated using the 10x Genomics 3′ Gene Expression v3.1 kit according to the manufacturer’s protocol. Sequencing was performed on the Illumina NovaSeq 6000 platform, targeting a depth of >50,000 reads per cell. Raw sequencing data were processed using the Cell Ranger pipeline (10x Genomics, v6.1) with default parameters. This included read trimming, alignment to the GRCh38/hg38 human genome reference, and unique molecular identifier (UMI) quantification. Cells with >20% mitochondrial gene expression or low transcript complexity (Genes per UMI < 0.8 for iTME cells and Genes per UMI < 0.5 for plasma cells) were excluded during quality control. Analysis herein focused exclusively on the CD138+ plasma cell compartment from the pre-treatment samples. Downstream processing and normalization were performed using the Seurat R package (v4.0). Batch effects across patients were addressed using canonical correlation analysis (CCA) for dataset integration. Plasma cell identity was confirmed by comparing transcriptional signatures with reference datasets, including the Cancer Cell Atlas and published immune cell single-cell transcriptomic studies. After quality control for feature counts and mitochondrial gene expression, there were a total of 48,150 plasma cells included.

### Single-Cell RNA Sequencing and Analysis (Baseline non-Plasma Cell Compartment)

For characterization of the immune microenvironment, the CD138- cell fraction from freshly isolated bone marrow aspirates was processed in parallel to the tumor-enriched CD138+ fraction. Raw FASTQ files were processed through the Cell Ranger (v6.1) pipeline using default parameters, including demultiplexing, alignment to the GRCh38/hg38 reference genome, and UMI-based quantification. Quality control was performed using Seurat (v4.0). Cells with >20% mitochondrial gene expression, <200 detected genes, or poor complexity (genes per UMI < 0.8 for immune/TME-associated cells) were removed. To ensure consistent annotation across datasets originating from different centers, samples from Moffitt Cancer Center and previously published Dhodapkar et al. BCD 2022 datasets were jointly integrated. Batch-effect correction was implemented using Harmony, which effectively aligned shared biological structure while preserving true inter-patient variability. Dataset integration, dimensionality reduction, clustering, and visualization were conducted using standard Seurat workflows (NormalizeData, FindVariableFeatures, ScaleData, and RunUMAP). Cell-type annotation was performed using a combination of unsupervised clustering and supervised reference mapping. Automated cell-type prediction was obtained using Azimuth (Human Bone Marrow reference). After QC filtering and batch integration, a total of 42,957 high-quality immune cells were retained for downstream analyses (substituting the exact number once finalized).

### Human myeloma cell line validations

K562 cells were stably transduced to co-express GFP and either wild-type BCMA or the P33S point mutation. Stable surface expression of BCMA was confirmed by flow cytometry using a fluorescently labeled anti-BCMA antibody.

Primary human T cells were isolated from healthy donor whole blood and stimulated using T Cell TransAct™ (Miltenyi Biotec) in the presence of IL-2 (200 IU/mL), with or without IL-7 (5 ng/mL) and IL-15 (10 ng/mL). On day 2 post-activation, T cells were transduced via spinoculation with lentiviral constructs encoding Ciltacabtagene autoleucel (Cilta-cel), Orvacabtagene autoleucel (Orva-cel), Idecabtagene vicleucel (Ide-cel), or a non-binding control CAR containing a dsRed reporter. CAR expression was confirmed and normalized based on dsRed fluorescence.

CART cells were co-cultured at a 1:1 ratio with K562-GFP target cells expressing each BCMA variant in RPMI 1640 medium supplemented with 10% fetal bovine serum. Target cell viability was monitored using the Incucyte® Live-Cell Analysis System (Sartorius) with GFP+ cell counts recorded every 2 hours for 96 hours.

### VkMYC mouse models

The generation and characterization of Vk*MYC transplantable lines have been reported elsewhere.^67^ One million cryopreserved splenocytes previously collected from tumor-bearing mice were transplanted by tail vein injection into wild-type C57BL/6 recipient mice. Upon tumor engraftment, detected as M-spike by serum protein electrophoresis (SPEP), mice received two weekly doses of BCMA-TCE at 1 mg/kg as previously described,^50^ and response to treatment was subsequently monitored by SPEP. Response was defined as >50% reduction in M-spike compared to day 0 levels. All genomic data derived from the Vk*MYC mouse were previously published and publicly available with accession code GSE255233.

### Genetically engineered mouse models

BIcγ1 mice were generated by crossing B6(Cg)-Gt(ROSA)26Sortm4(Ikbkb)Rsky/J mice with constitutively active NF-κB signaling by Ikbkb expression and GFP expression, B6.Cg-Tg(BCL2)22Wehi/J mice with constitutive BCL2 expression, and cγ1-cre mice (B6.129P2(Cg)-Ighg1tm1(cre)Cgn/J). Tp53-BIcγ1 and NSD2-BIcγ1 mice were generated by crossing BIcγ1 mice with B6.129P2-Trp53tm1Brn/J mice with Trp53 deletion, or with Rosa26-hMMSET-IIStop-floxed mice carrying the human MMSET-II gene sequence, which was maintained in heterozygosity. MIcγ1 mice were generated by crossing B6(Cg)-Gt(ROSA)26Sortm4(Ikbkb)Rsky/J mice and C57BL/6N-Gt(ROSA)26Sortm13(CAG-MYC,-CD2*)Rsky/J mice with c-MYC expression, and cγ1-cre mice. All models have been previously reported,^46^now backcrossed >10 generations to obtain a pure C57BL/6 background. To induce the formation of GFP+ transgenic plasma cells in mice housed under specific pathogen-free (SPF) conditions, animals were subjected to T cell-mediated immunization with sheep red blood cells administered intraperitoneally from 8 weeks of age. In all experiments, age-matched cγ1-cre mice similarly immunized were used. BIcγ1, Trp53-BIcγ1, NSD2-BIcγ1 and MIcγ1 mice progressively developed MM in the BM and died of the disease around 10, 8.5, 10, and 7 months of age, respectively. Bulk and single-cell RNA sequencing were applied to GFP+CD138+B220- MM cells isolated from the BM of the different mouse strains at the time of death, following reported methods.^46^ All animals were maintained under SPF conditions at the animal facility of the University of Navarra. The study was approved by the Ethical Committee of Animal Experimentation of the University of Navarra and the Health Department of the Navarra Government.

### Preclinical therapy trial testing responses to BcmaxCd3

A murine BcmaxCd3 surrogate bispecific antibody was administered to Trp53-BIcγ1 mice with heterozygous deletion of Trp53 (n=12) and to MIcγ1 mice with wild-type Trp53 (n=13), while control arms received isotype antibody (n=10 and n=7, respectively). Before therapy initiation, tumor burden was estimated in each mouse by measuring the immunoglobulin gamma fraction (M spikes) in serum by electrophoresis. Once clonal M spikes were detected, BcmaxCd3 or isotype at 1 mg/kg were administered by intraperitoneal injection once weekly for 8 weeks. Therapy responses were determined by comparing serum M spikes before therapy and then every 4 weeks. Mice were then followed up until death. Survival was estimated by Kaplan-Meier survival curves in the different treatment cohorts.

### Statistical analysis

The study sample size and experiments were based on sample availability. Statistical comparisons were conducted using Log Rank, Wilcoxon and Fisher’s exact tests, depending on whether the analysis involved continuous or categorical variables. All reported p-values are two-sided unless stated otherwise. The Kaplan–Meier method was employed to estimate time-to-event distributions. PFS was defined as the time from treatment initiation to either disease progression. All visualizations and figures were created using R-Studio Version 2023.09.1+494, Adobe Illustrator, and Biorender.

## DATA AVAILABILITY

All sequencing data are available are in the process of being uploaded on EGA

## ACKNOWLEDGMENTS.

This work was supported by the ASH Scholar, Leukemia and Lymphoma Society (LLS; 6677-24), Myeloma Solutions Fund (MSF), The Paula and Rodger Riney Foundation, Sylvester Comprehensive Cancer Center NCI Core Grant (P30 CA 240139), Memorial Sloan Kettering Cancer Center National Cancer Institute (NCI) Core Grant (P30 CA 008748), and the Hematology Oncology Pharmacy Association (HOPA).

CLF: This study was supported (in part) by institutional research funding from Bristol Meyers Squibb in addition to research funding by the Pentecost Family Myeloma Research Center. This work was also supported by the National Cancer Institute (P30CA076292, PI Cleveland).

F.L.L. is supported in part by the Leukemia and Lymphoma Society as a Clinical Scholar, the National Cancer Institute (R01CA244328-03), and generous donations from the Hano, Hyer family, Hoenle Family and the Thiel family.

Moffitt team: The authors sincerely thank our multiple myeloma patients and their families for donating their samples for research purposes. This research was made possible through the Total Cancer Care (TCC™) protocol at the H. Lee Moffitt Cancer Center & Research Institute, an NCI-designated Comprehensive Cancer Center (P30-CA076292). We are also grateful for the expert assistance and service of the Tissue Core, Molecular Genomics, Cancer Pharmacokinetics and Pharmacodynamics, Flow Cytometry, and Biostatistics and Bioinformatics Cores of Moffitt Cancer Center. Additionally, they would like to acknowledge the ongoing collaboration with Aster Insights for aspects of the molecular analysis (bulk whole exome sequencing) as well as bioinformatic analysis. This work was supported by invaluable philanthropic support from the Pentecost Family Myeloma Research Center (to R.B., K.H.S., A.S., M.A.) and Bowlin Family Foundation (K.H.S).

ML and JAM-C are supported by the International Myeloma Society (IMS), the Spanish Ministry of Health (ISCIII-FIS and CIBERONC-CB16/12/00489), and Fundacion Arnal Planelles

R.B.: Janssen, BMS, AbbVie, Regeneron, Karyopharm

FM is supported by Leukemia & Lymphoma Society (6677-24), by the International Myeloma Society (IMS), ASH Scholar, Department of Defense (DoD), and NIH-NCI.

KHM has received funding from the Multiple Myeloma Research Foundation, the American Society of Hematology, and the International Myeloma Society.

TJ is supported by EHA Physician Scientist Research Grant RG-202409-06772. The work was supported by OPJAK SALVAGE project (CZ.02.01.01/00/22_008/0004644).

## AUTHOR CONTRIBUTIONS STATEMENT

F.M. and C.F., designed and supervised the study, collected, generated, and analyzed the data and wrote the paper. S.U., F.L., I.A., and K.S., designed and supervised the study, collected the samples and wrote the paper. O.L.., designed and supervised the study and wrote the paper. P.N.: collected the samples and analyzed the data. K.M collected the samples, generated, and analyzed the data, and wrote the paper. M.La. generated and analyzed the data related to the Trp53 mouse model, and write the paper; B.Z., H.L., M.D., R.S., M.M., B.Z., M.P., A.S.S., P.R.S., N.L., S.A., E.F., E.B., P.A., D.K.H., X.S., X.Z., X.Z., M.B.M., J.T., J.J.C., analyzed the data, R.B., M.A., A.R., S.M., S.G., collected the samples and clinical data. P.H. provided the murine surrogate BcmaxCd3 T cell engager for the MIcγ1 and Tp53-BIcγ1 mice. L.B., M.C., J.A.M.-C. supervised and generated the mouse data.

## COMPETING INTERESTS STATEMENT

O.L. has received research funding from: National Institutes of Health (NIH), National Cancer Institute (NCI), U.S. Food and Drug Administration (FDA), Multiple Myeloma Research Foundation (MMRF), International Myeloma Foundation (IMF), Leukemia and Lymphoma Society (LLS), Myeloma Solutions Fund (MSF), Paula and Rodger Riney Multiple Myeloma Research Program Fund, the Tow Foundation, Perelman Family Foundation, Rising Tide Foundation, Amgen, Celgene, Janssen, Takeda, Glenmark, Seattle Genetics, Karyopharm; Honoraria/ad boards: Adaptive, Amgen, Binding Site, BMS, Celgene, Cellectis, Glenmark, Janssen, Juno, Pfizer; and serves on Independent Data Monitoring Committees (IDMCs) for clinical trials lead by Takeda, Merck, Janssen, Theradex.

K.H.M. has received funding from the Multiple Myeloma Research Foundation, The American Society of Hematology and the International Myeloma Society.

F.M.: consulting for Medidata and Sanofi.

B.D. reports consulting via independent data review committee for Janssen.

J.A.M.-C. has received research funding from Roche-Genentech, Bristol Myers Squibb, Janssen, Regeneron, Priothera Pharmaceuticals, Palleon Pharmaceuticals, AstraZeneca, and K36 Therapeutics outside of this work, is founder and holds stock options of MIMO Biosciences, and declares a patent licensed to MIMO Biosciences on the generation, knowhow and use of the MM models described here.

A.S.S reports research funding to the institution from AbbVie and Karyopharm, outside the submitted work.

K.H.S. reports honoraria from Amgen, Bristol Myers Squibb, Janssen, Adaptive, Sanofi, Regeneron, and Takeda, and research funding to the institution from AbbVie, Karyopharm, and Pfizer outside the submitted work.

R.B. reports research support: Janssen, BMS, Abbvie, Regeneron, Karyopharm

D.K.H reports research funding from Bristol-Myers Squibb, Janssen, Karyopharm, Kite Pharma, and Adaptive Biotech; Consulting or advisory role for Bristol Myers Squibb, Janssen, Legend Biotech, Pfizer, Kite Pharma/Gilead Sciences, AstraZeneca, and Karyopharm

F.L.: Scientific Advisory Role/Consulting Fees: A2, Adaptive Biotechnologies, Adaptimmune, Allogene, Amgen, Astra-Zeneca, Bluebird Bio, BMS, Calibr, Caribou, EcoR1, Gerson Lehrman Group (GLG), Iovance, Kite Pharma, Janssen, Legend Biotech, Miltenyi, Novartis, Sana, Pfizer, Poseida Data Safety Monitoring Board: Data and Safety Monitoring Board for the NCI Safety Oversight CAR T-cell Therapies Committee. Research Contracts or Grants to my Institution for Service: 2SeventyBio (Institutional), Allogene (Institutional), BMS (Institutional), Incyte (Institutional), Kite Pharma (Institutional), Leukemia and Lymphoma Society Scholar in Clinical Research (PI: Locke), Mark Foundation, National Cancer Institute, Patents, Royalties, Other Intellectual Property: Several patents held by the institution in my name (unlicensed) in the field of cellular immunotherapy. Education or Editorial Activity: Aptitude Health, ASH, ASTCT, Clinical Care Options Oncology, Society for Immunotherapy of Cancer CLF Honoraria/consulting BMS/Celgene, ONK therapeutics & Janssen; research funding from BMS, Janssen and Roche/Genentech.

All other authors have no conflicts of interest to declare.

