## Supplementary Figures for "Plasma cell identity escape drives resistance to anti-BCMA T-cell–redirecting therapy in multiple myeloma"

**Supplementary Figure 1. A-B)** Kaplan Meier showing clinical impact in term of PFS of EMD (**A**) and prior anti-BCMA therapy (**B**) in patients treated with CART or T-cell engagers (TCE). **C-D**) Impact of soluble BCMA (sBCMA) (**C**), and myCARe score (**D**) on PFS in patients treated with CART. **E-F)** Kaplan Meier showing PFS in patients treated with Ciltacel vs idecel (**E**) and Elranatanab vs Teclistamab (**F**).

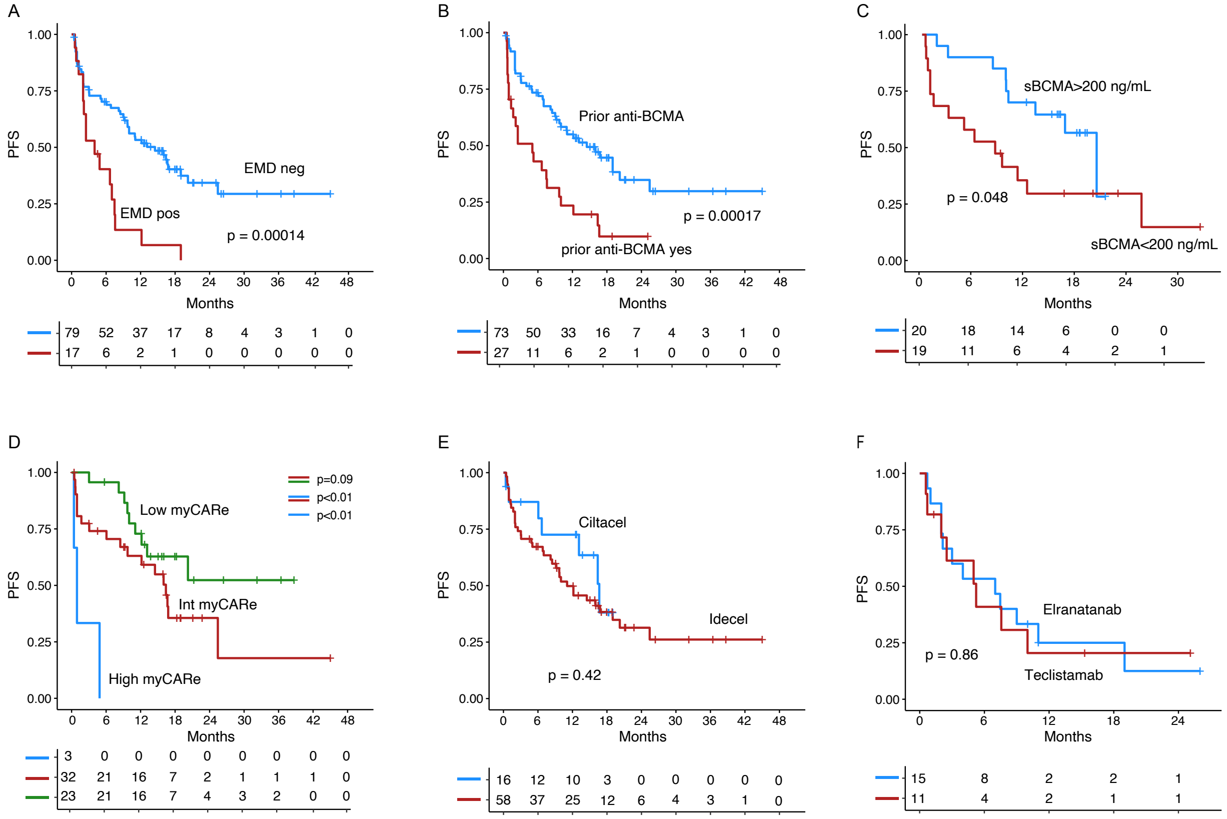

**Supplementary Figure 2. CAR T cell cytotoxic potential is not impacted by P33S point mutation.** Various publicly available BCMA-targeted CAR sequences were used to engineer CAR T cells. Anti-tumor efficacy was evaluated by live cell imaging of K562 (GFP+) cells engineered to express either (A) wild type BCMA or (B) BCMA with P33S mutation. (Representative data shown from one of two replicates using separate human donor T cells).

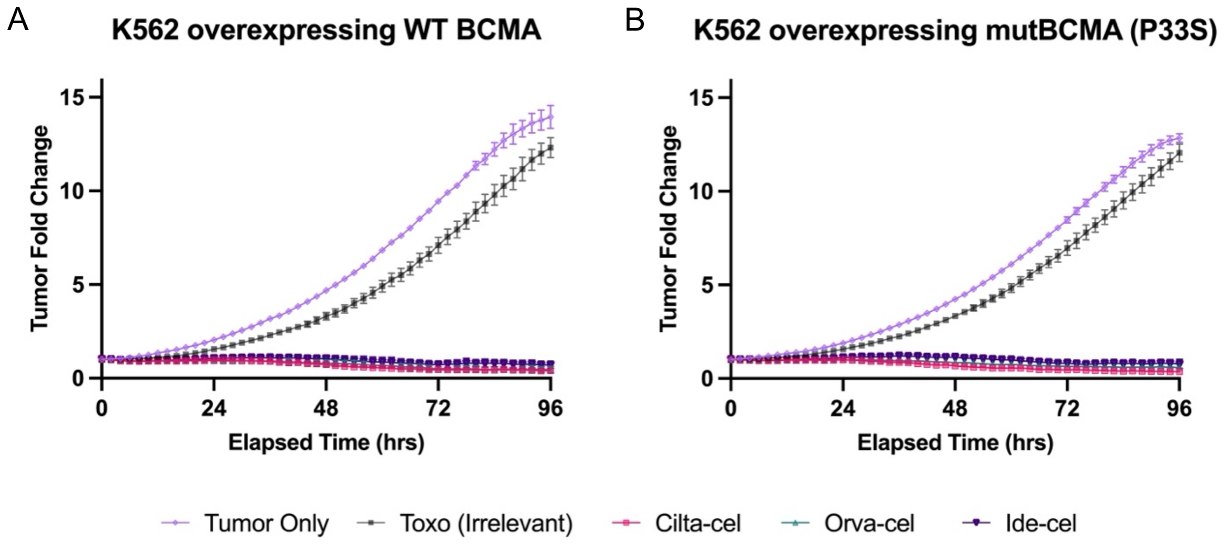

**Supplementary Figure 3.** Phylogenetic tree reconstruction for patients with one or more samples collected after anti-BCMA CART

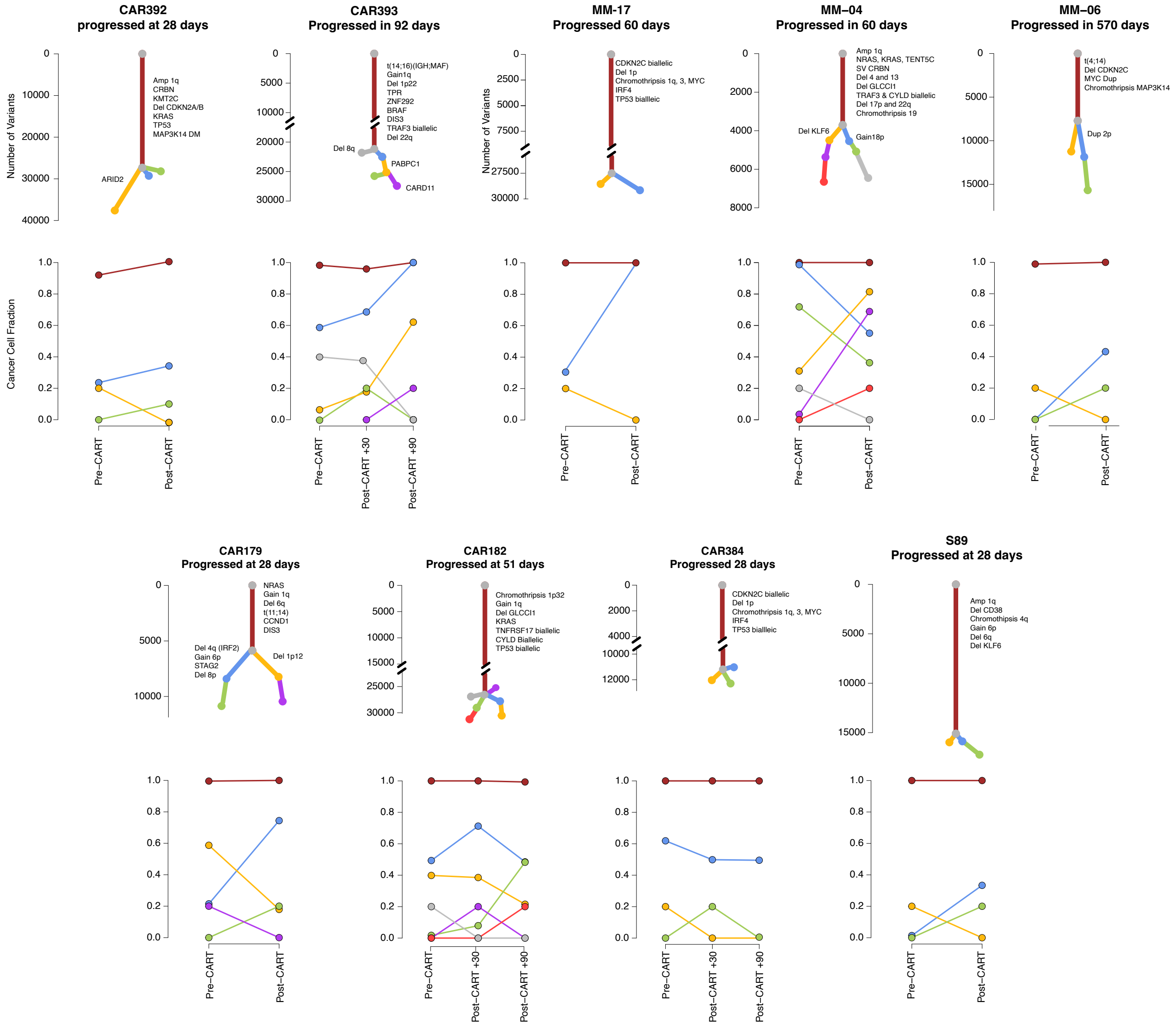

**Supplementary Figure 4. Absence of immune exhaustion/activation in the T CD4 of refractory (REF) patients compared to durable responders (DR).**

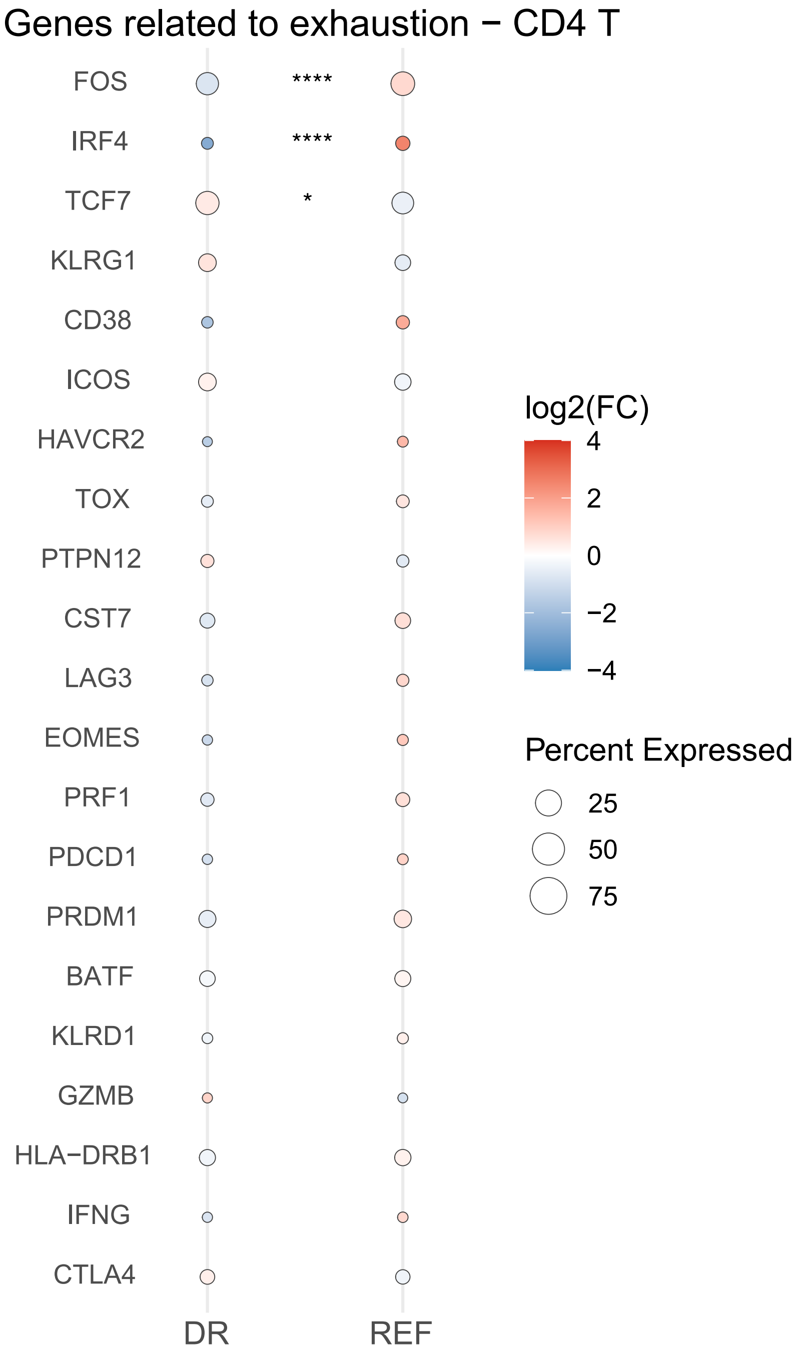

**Supplementary Figure 5. A-B)** Impact of CD38 loss in patients treated with T-cell engagers (TCE; **A**) and CART (**B**). **C-D**) Correlation between *CD38* and *TNFSRF17* expression across 683 RRMM (**C**) and 1036 NDMM (**D**) with available RNAseq, respectively. **E)** Kaplan-Meier showing the impact of *XBP1* loss before treatment with anti-BCMA CART in term of PFS. **F-G)** Correlation between *CD38* and *TNFSRF17* expression across 683 RRMM (**F**) and 1036 NDMM (**G**) with available RNAseq, respectively. **H-I**) Impact of CD38 loss in patients treated with TCE (**H**) and CART (**I**).

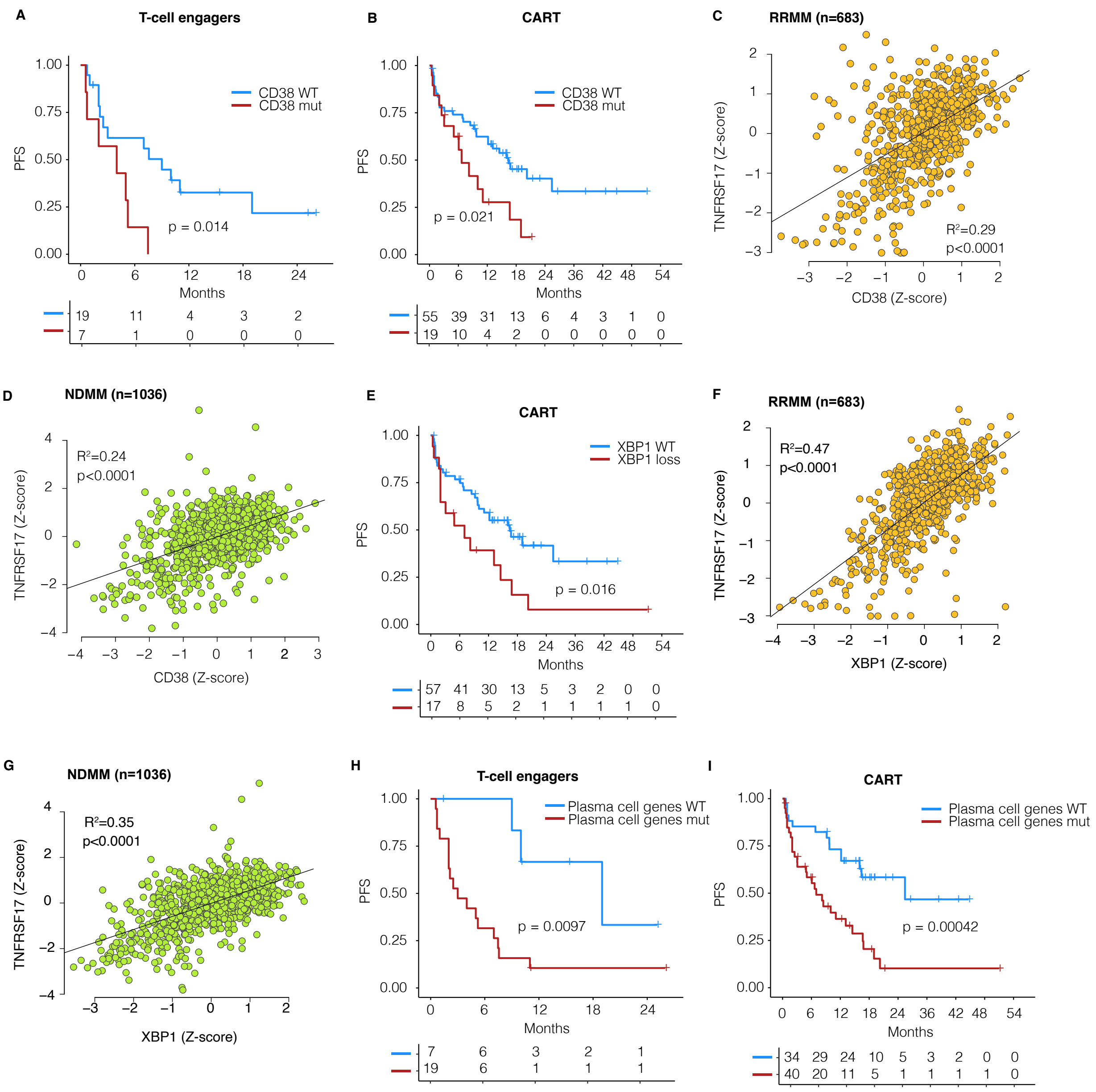

**Supplementary Figures 6. A-B**) Impact of 1q gain/amplification  in patients treated with CART (**A**) and TCE (**B**). **C-D**) Impact of TP53 mutation in patients treated with CART (**C**) and TCE (**D**). **E)** Correlation between *TP53* and *TNFSRF17* expression across 1036 NDMM.

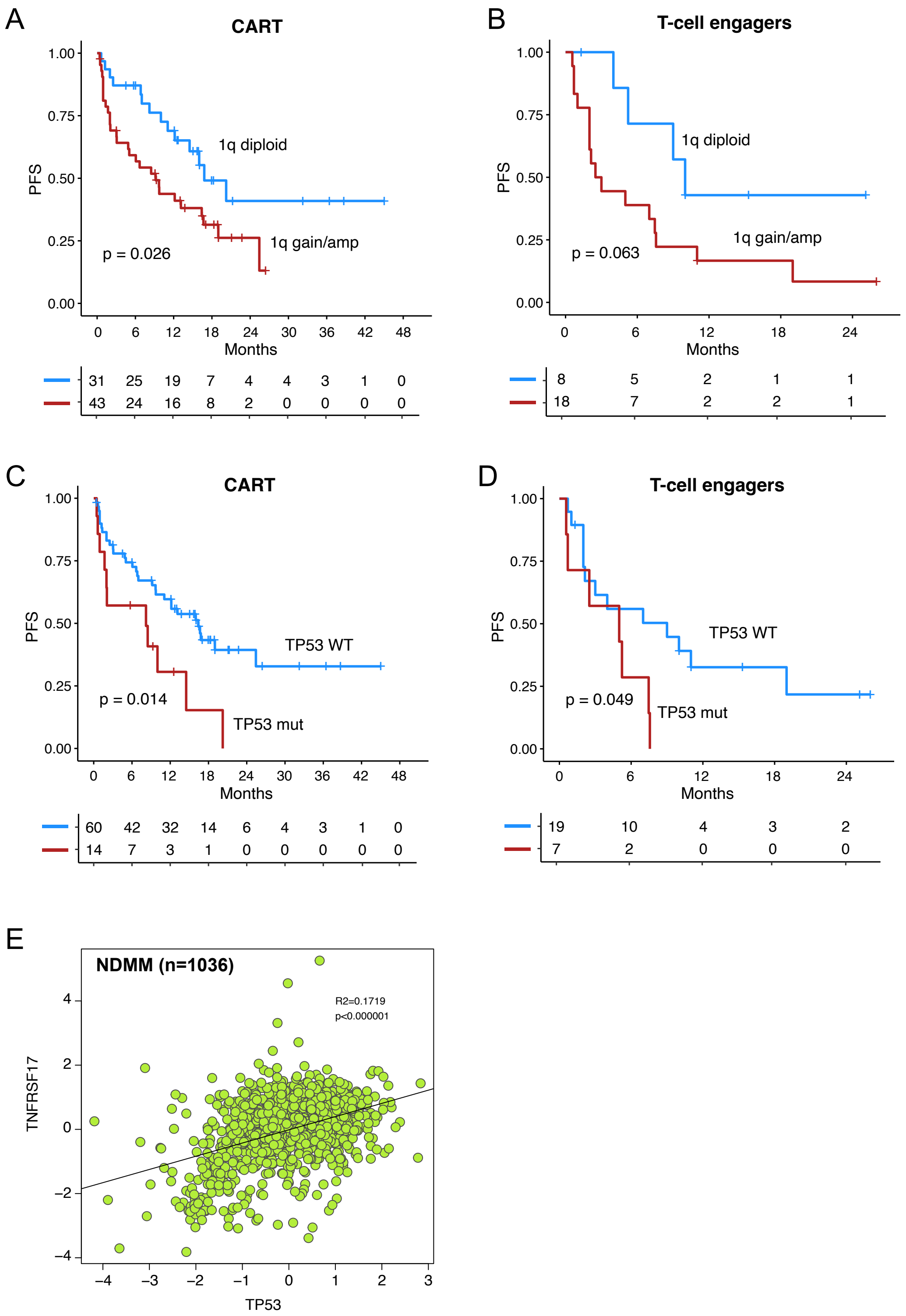

**Supplementary Figure 7. A)** Chance in gamma fraction between Tp53BIcγ1 and MIcγ1 mice after treatment with anti-BCMA TCE. **B-C)** M-spike levels (serum gamma/albumin) measured over time in C57BL/6 wild type mice engrafted with Vk35633 (**E**) and Vk27181 (**F**), myeloma cells after treatment with two weekly doses (1 mg/kg) of a murine anti-BCMA TCE. Each line represents an individual mouse.

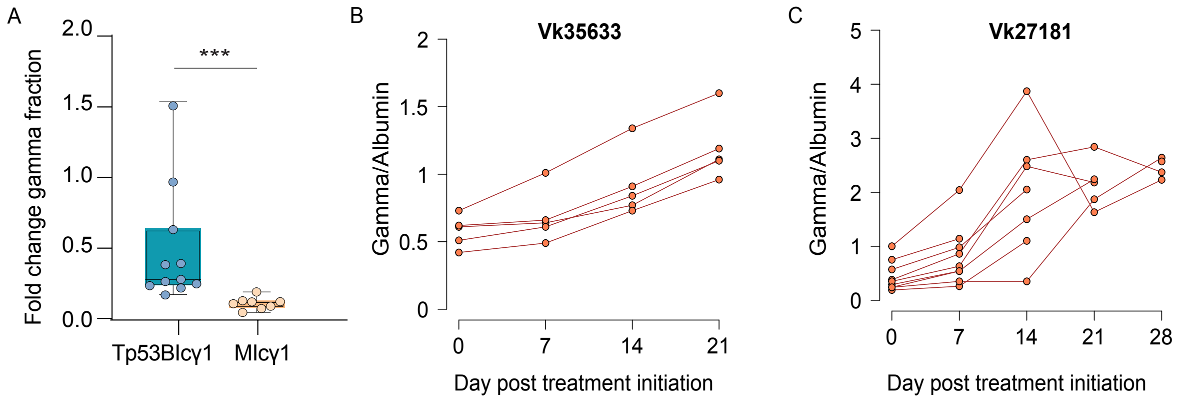
